# Unravelling the regulation pathway of photosynthetic AB-GAPDH

**DOI:** 10.1101/2021.11.21.469434

**Authors:** Roberto Marotta, Alessandra Del Giudice, Libero Gurrieri, Silvia Fanti, Paolo Swuec, Luciano Galantini, Giuseppe Falini, Paolo Trost, Simona Fermani, Francesca Sparla

**Affiliations:** Electron Microscopy Facility (EMF), Italian Institute of Technology (IIT), 16163 Genova, Italy; Department of Chemistry, University of Rome Sapienza, 00185 Rome, Italy; Department of Pharmacy and Biotechnology–FaBiT, University of Bologna, 40126 Bologna, Italy; Department of Chemistry G. Ciamician, University of Bologna, 40126 Bologna, Italy; Biosciences Department, University of Milan, 20133 Milan, Italy; Interdepartmental Centre for Industrial Research Health Sciences & Technologies, University of Bologna, 40064 Bologna, Italy

**Keywords:** Photosynthesis, Calvin-Benson cycle, Redox regulation, Glyceraldehyde-3-phosphate dehydrogenase, Cryo-electron microscopy, Small angle X-ray scattering

## Abstract

Oxygenic phototrophs perform carbon fixation through the Calvin–Benson cycle. Different mechanisms adjust the cycle and the light-harvesting reactions to rapid environmental changes. Photosynthetic glyceraldehyde 3-phosphate dehydrogenase (GAPDH) is a key enzyme of the cycle. In land plants, different photosynthetic GAPDHs exist: the most abundant formed by hetero-tetramers of A and B-subunits, and the homo-tetramer A_4_. Regardless of the subunit composition, GAPDH is the major consumer of photosynthetic NADPH and for this reason is strictly regulated. While A_4_-GAPDH is regulated by CP12, AB-GAPDH is autonomously regulated through the C-terminal extension (CTE) of B-subunits. Reversible inactivation of AB-GAPDH occurs via oxidation of a cysteine pair located in the CTE, and substitution of NADP(H) with NAD(H) in the cofactor binding domain. These combined conditions lead to a change in the oligomerization state and enzyme inactivation. SEC-SAXS and single-particle cryoEM analysis disclosed the structural basis of this regulatory mechanism. Both approaches revealed that (A_2_B_2_)_n_-GAPDH oligomers with n=1, 2, 4 and 5 co-exist in a dynamic system. B-subunits mediate the contacts between adjacent A_2_B_2_ tetramers in A_4_B_4_ and A_8_B_8_ oligomers. The CTE of each B-subunit penetrates into the active site of a B-subunit of the adjacent tetramer, while the CTE of this subunit moves in the opposite direction, effectively preventing the binding of the substrate 1,3-bisphosphoglycerate in the B-subunits. The whole mechanism is made possible, and eventually controlled, by pyridine nucleotides. In fact, NAD(H) by removing NADP(H) from A-subunits allows the entrance of the CTE in B-subunits active sites and hence inactive oligomer stabilization.

**Significance Statement:** In land plants, glyceraldehyde 3-phosphate dehydrogenase (GAPDH) unique sink of reducing power of the entire Calvin-Benson cycle, is finely regulated. Based on the redox state and substrates concentration, its heteromeric form AB-GAPDH oscillates between a fully active heterotetramer (A_2_B_2_) and inactive oligomers. Experimental evidence demonstrates that GAPDH inactivation depends on the formation of dimers, tetramers or pentamers of A_2_B_2_-modules, linked together by C-terminal extensions (CTE) of B-subunits that extrude from one modular tetramer and occupy two active sites of the adjacent one. This molecular mechanism along with the unexpected observed dynamism of the system, shed light on how the Calvin-Benson cycle is modulated in function of the light environmental changes.

## Introduction

Oxygenic photosynthesis sustains all life on Earth reducing carbon dioxide into carbohydrates while photo-oxidizing water into oxygen. The photosynthetic electron transport chain, strictly dependent on light, provides energy (ATP) and reducing power (NADPH) for the metabolic phase of the process. By consuming ATP and NADPH, carbohydrates are produced from CO_2_ by the Calvin-Benson cycle (1–3). Despite the historical distinction between the two phases of photosynthesis, the entire process occurs during the day through a complex and diversified regulatory system that harmonizes the rate of carbon fixation with the rate of conversion of light energy into chemical energy (4–6).

Thioredoxins (TRXs) represent one of the wake-up calls of the Calvin-Benson cycle at dawn. Through the TRX/ferredoxin system part of the reducing power originated by the photosystem I induces the activation of the cycle in a TRX dependent manner (7–9). In land plants, phosphoribulokinase (PRK) (10–12), fructose 1,6-bisphosphatase (FBPase) (13, 14), sedoheptulose-1,7-bisphosphatase (SBPase) (14) and the AB-isoform of glyceraldehyde 3-phosphate dehydrogenase (GAPDH) are direct target of TRXs(15, 16).

GAPDH catalyzes the only reducing step of the Calvin-Benson cycle and is the major consumer of the photosynthetically produced NADPH. Two isoforms of photosynthetic GAPDH coexist in the chloroplast stroma of land plants: a homo-tetramer exclusively made of A subunits, and a hetero-tetramer containing both A and B-subunits (3, 17) that can form higher order oligomers (18, 19). The structure of A_4_- and A_2_B_2_-GAPDH is similar and highly conserved among GAPDHs (20, 21). Although the regulation of both isoforms occurs by interaction with CP12 and PRK, AB-GAPDH shows an additional autonomous regulation (3, 22).

CP12 is a small conditionally disordered protein containing two pairs of conserved cysteines (23, 24). The C-terminal pair, with a midpoint redox potential (*E*_m,7.9_) of -352 mV, binds GAPDH, while the less negative potential N-terminal disulfide (*E*_m,7.9_ = -326 mV) recruits PRK into the complex (3, 25). Recently, the structure of A-GAPDH/CP12/PRK complex has been solved, enlightening the molecular mechanisms involved in complex formation and redox regulation (12, 26, 27).

AB-GAPDH performs the CP12-independent regulation through the presence of a 30 amino acid tail specific of the B-subunit (3, 15, 21, 28). This C-terminal extension (CTE) is highly homologous to the C-terminal region of CP12 and it has been postulated that the B-subunit results from the fusion between the A-subunit and the C-terminal half of CP12 (3, 29–31). Like CP12, also the CTE contains two cysteines that can be possibly engaged in a disulfide bridge.

AB-GAPDH exhibits its own propensity to vary the oligomeric state from active heterotetramers to inactive hexadecamers (18, 28, 32, 33). The transition between the oligomeric states depends not only on the redox state of the CTE, but also on the type of cofactor (NADP(H) or NAD(H)) and on the substrate 1,3-bisphosphoglycerate (BPGA) availability (15, 34).

With the aim of disclosing the molecular mechanism that drives the oligomerization of AB-GAPDH, here we report a multi-approach structural study of the AB-GAPDH system by small angle X-ray scattering coupled with size exclusion chromatography (SEC-SAXS) and single-particle cryoelectron microscopy (cryoEM). Both experimental approaches highlight an unexpected dynamism of AB-GAPDH. Moreover, cryoEM reveals that pairs of B-subunits belonging to adjacent tetramers, mutually exchange their CTEs. Protruding like hooks, CTEs dock and penetrate in the active site of the adjacent tetramer blocking the access of the substrate.

## Results and Discussion

### Fingerprinting multiple oligomeric states of AB-GAPDH with SEC-SAXS

SEC-SAXS data were collected on active (NADP^+^-bound) and inactive (NAD^+^-bound) AB-GAPDH oligomers previously analyzed by Dynamic Light Scattering (DLS). Average hydrodynamic radius (R_h_) values of 52 and 100 Å corresponding to apparent molecular weight (MW) of 159 and 736 kDa, were obtained for active and inactive forms, respectively. As a reference, the theoretical MW of A_2_B_2_-GAPDH tetramers is 149 kDa. SAXS experiments showed that all samples presented a systematic variation of dimensional parameters, underlying the presence of different oligomers in addition to the more abundant A_2_B_2_ and A_8_B_8_ species expected in active and inactive samples, respectively (Fig. 1*A* and *SI Appendix*, Fig. S1) (15, 17, 21, 28). Statistically superimposable frames showing a constant gyration radius (R_g_) were identified and averaged to obtain representative SAXS profiles (Fig. 1*A* and *SI Appendix*, Table S1) interpretable as AB-GAPDH oligomers on the basis of their dimensional parameters and distance distribution functions (P(r)) (Fig. 1 and *SI Appendix*, Fig. S1 and Table S2).

**Figure 1.**
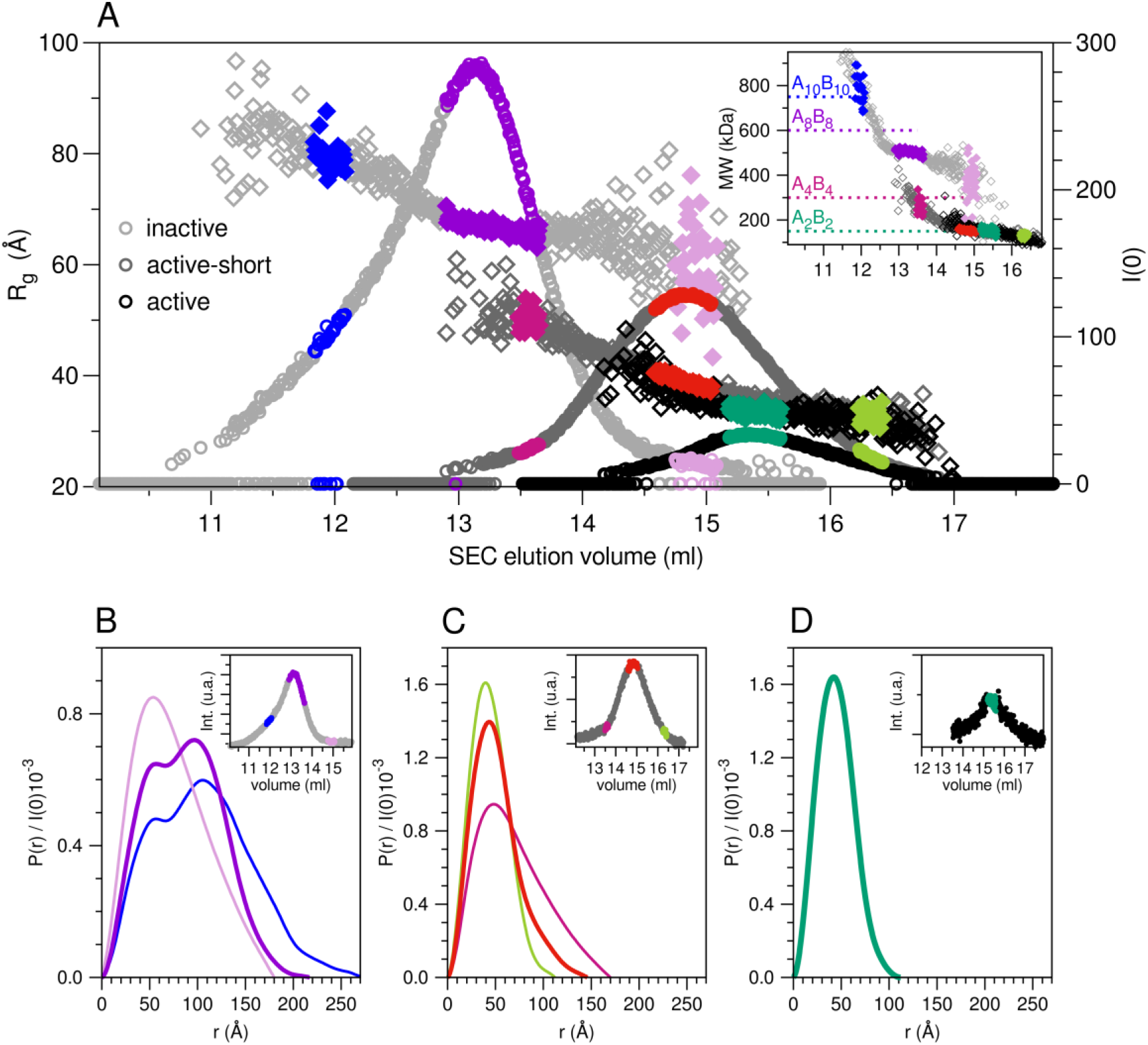
SEC-SAXS elution profiles and P(r) functions of AB-GAPDH species. (A) Parameters derived from the analysis of SAXS frames for the three AB-GAPDH samples: inactive (light grey symbols, maximum at 13 ml), active-short (grey symbols, maximum at 14.8 ml) and active (black symbols, maximum at 15.4 ml), are shown as a function of the SEC elution volume. The datapoints belonging to the frames averaged to obtain the selected scattering profiles are highlighted with a colour code. The elution profile given by the scattering intensity at zero angle (I(0), circles, right ordinate axis) is plotted together with the radius of gyration (R_g_) obtained from the Guinier approximation (diamonds, left ordinate axis). In the inset, molecular weight (MW) estimated from the Porod volume (MW(V_P_), diamonds). The MWs expected on the basis of the protein sequence for (A_2_B_2_)_n_ oligomers with n=1, 2, 4 and 5 are reported as dashed lines for reference. P(r) functions calculated from the selected scattering profiles in the elution of the samples: (B) inactive, (C) active-short, (D) active; in the insets the elution profiles given by the SAXS integrated intensity are also shown.

In the inactive sample, the predominant species showed a R_g_ of 67 Å, a maximum size (D_max_) of 200 Å and a MW between 500 and 600 kDa, compatible with the expected A_8_B_8_ oligomer (*SI Appendix*, Table S1). In addition, a larger construct was identified, with a R_g_ around 80 Å, a D_max_ of 280 Å and an estimated MW between 650 and 700 kDa, suggesting an A_10_B_10_ stoichiometry. A less abundant and smaller component was also observed at larger elution volumes (Fig. 1*A*, pink symbols). The estimation of its R_g_ and MW was more uncertain. The related P(r) profile showed a D_max_ around 150 Å and only one maximum around 50 Å, clearly distinguishable from the bimodal P(r) function of A_8_B_8_ (Fig. 1*B*). A similar P(r) profile (Fig. 1*C*) was calculated also at the beginning of the elution of the active-short sample again showing a wide range of estimated MWs (Fig. 1*A* inset, purple diamonds). This sample was obtained from 2 hours incubation of the inactive sample under activating conditions (see *SI Appendix*, Materials and Methods). On this basis, the co-existence of species with intermediate stoichiometries possibly centered on A_4_B_4_ and being in exchange equilibria with higher oligomers (inactive sample) or smaller oligomer (active-short sample) can be envisioned. The presence *in vivo* of the A_4_B_4_ was already reported in different plant species (33, 35, 36) besides the common A_2_B_2_ and A_8_B_8_-GAPDH forms, supporting the idea that this oligomer is not only an intermediate in the aggregation of A_2_B_2_ to A_8_B_8_, but even an essential player for AB-GAPDH regulation.

The R_g_ and D_max_ (Fig. 1A, red symbols, *SI Appendix*, Fig. S1 and Table S2) detected at the elution maximum of the active-short sample (14.8 ml) would suggest a fast-exchange dynamic equilibrium between A_2_B_2_ and higher order oligomers (possibly involving A_4_B_4_ as an intermediate species) in the elution conditions, giving rise to the impossibility to detect distinct SEC peaks (37).

The dimensional parameters of the active-short sample decreased gradually towards larger retention volumes and at the end of the elution, the structural parameters agreed with those found at the elution maximum of the active sample, i.e. a R_g_ of 34 Å and a D_max_ around 100 Å, compatible with an A_2_B_2_ tetramer (Fig. 1*D*).

### Single-particle cryoEM analysis confirms the heterogeneity of inactive AB-GAPDH

In agreement with SAXS results, in inactivating conditions the single-particle analysis revealed the coexistence of different oligomeric states of the enzyme (Fig. 2). Projections related to different GAPDH oligomers, namely A_2_B_2_, A_4_B_4_, A_8_B_8_ and A_10_B_10_, were clearly present in negative stain and cryoEM micrographs (*SI Appendix*, Fig. S2). They were also present in the 2D and 3D classifications performed on the complete GAPDH data set (Figs. 2*A* and *SI Appendix*, Fig. S3*A*).

**Figure 2.**
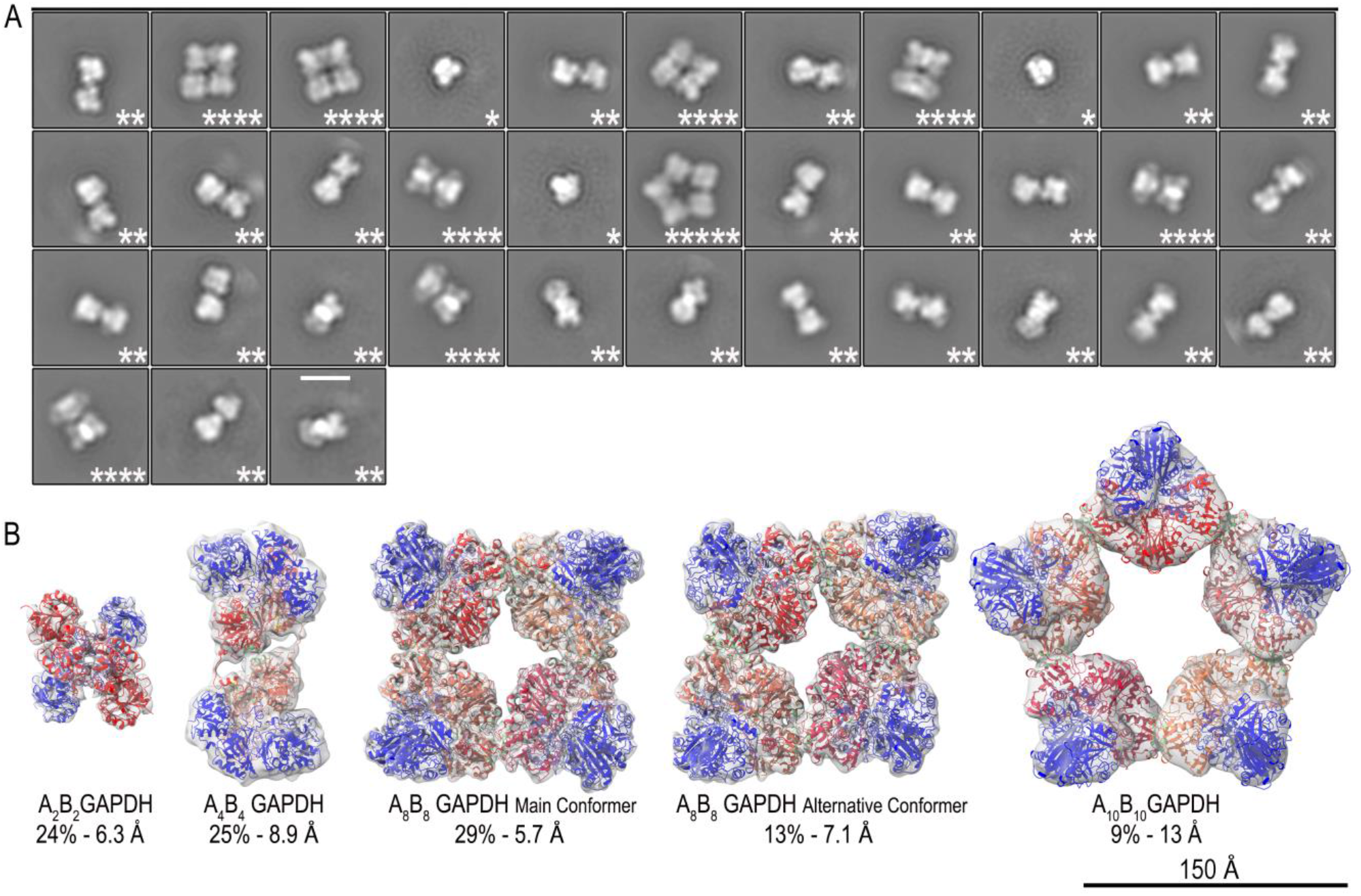
AB-GAPDH oligomers. (A) Representative single-particle 2D classification obtained from the complete GAPDH data set showing the presence of class averages attributable to A_2_B_2_, A_4_B_4_, A_8_B_8_ and A_10_B_10_ oligomers. For each species, the number of A_2_B_2_ tetramers is indicated by asterisks. The scale bar is 150 Å. (B) GAPDH oligomer cryoEM density maps fitted with models derived from the crystal structure of the oxidized A_2_B_2_ complexed with NADP+ (PDB ID code 2PKQ) (21). The O/Q, A/C, E/G, K/I and M/S B-subunits are represented in red, tomato, crimson, coral and indian red, respectively. The A-subunits are in blue. The numbers below the cryoEM electron density maps represent the oligomer relative abundances and their resolutions, respectively.

An estimation of the relative abundance of each oligomer obtained from the number of refined particles, showed that the A_8_B_8_ hexadecamer is the most abundant species (42%), albeit in two distinct conformers, named main (29%) and alternative (13%) (Fig. 2*B*). The A_4_B_4_ octamer (25%) and the A_2_B_2_ tetramer (24%) are less abundant. The remaining 9% corresponds to the A_10_B_10_ icosamer.

The cryoEM density map of the A_2_B_2_ tetramer was determined at 6.3 Å (Fig. 2*B* and *SI Appendix*, Fig.S4). Its superimposition to the crystal structure of oxidized A_2_B_2_-GAPDH (PDB ID 2PKQ) (21) does not point out significant conformational differences.

The 8.9 Å A_4_B_4_ cryoEM density map is a dimer with c1 symmetry formed by two A_2_B_2_ tetramers rotated each other by approximately 180° (Figs. 2*B* and 3*A* and *SI Appendix* Figs. S5*A, B*). Imposition of c2 symmetry in the 3D refinement process produced a less resolved reconstruction.

The A_8_B_8_ hexadecamer exists in two conformations, both with c2 symmetry and formed by two A_4_B_4_ dimers. The 5.7 Å cryoEM density map of the main conformer shows a central cavity with an area of about 1763 Å^2^ (Figs. 2*B* and 3*D*-*I* and *SI Appendix* Fig. S5*C*). Compared to the main conformer, the two A_4_B_4_ dimers of the alternative conformer (solved at 7.1 Å) are slightly shifted in the x direction, one in respect to the other, and the central cavity has a similar area (1738 Å^2^) (Fig. 2*B* and *SI Appendix*, Fig. S6*A*).

**Figure 3.**
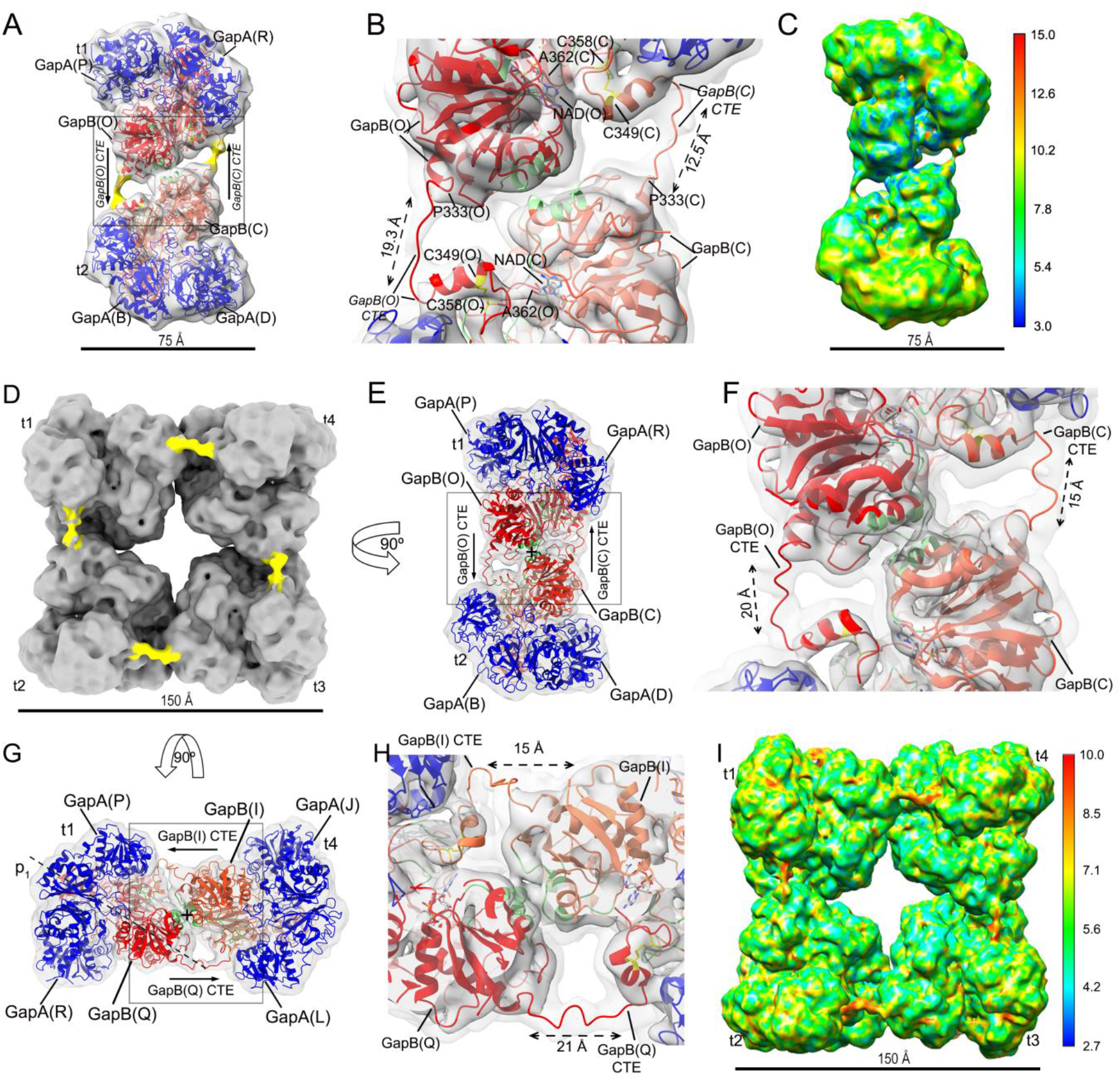
A_4_B_4_ and A_8_B_8_ oligomers. (A) CryoEM density map of the A_4_B_4_ oligomer at 8.9 Å resolution. The map, shown at low density threshold, reveals two regions (highlighted in yellow) connecting the t1 and t2 A_2_B_2_ tetramers. (B) Detail of the region boxed in (A). (C) CryoEM electron density map of the A_4_B_4_ oligomer filtered according to ResMap local resolution. (D) CryoEM electron density map of the A_8_B_8_ oligomer shown at a low density threshold. Note the connecting regions (highlighted in yellow) among the GAPDH tetramers t1-t4. (E) Side view of the maps in (D) showing the t1 and t2 tetramers. (F) Detail of the boxed region in (E). (G) Side view of the map in (D) showing the t1 and t4 tetramers. (H) Detail of the region boxed in (G). (I) CryoEM electron density map of the A_8_B_8_ oligomer filtered according to ResMap local resolution. All maps are fitted with their corresponding model derived from the crystal structure of the oxidized A_2_B_2_ complexed with NADP^+^ (PDB ID code 2PKQ) (18). The O/Q, A/C, E/G, K/I and M/S B-subunits are represented in red, tomato, crimson, coral and indian red, respectively. The A-subunits are in blue. In (B), (F) and (H), the densities of the 3D reconstructions are displayed at two different isosurface levels (higher in dark gray and lower in light gray) and the interfacing residues between adjacent GAPDH tetramers are highlighted in green.

Finally, the 13 Å A_10_B_10_ electron density map is a pentamer with c5 symmetry and a central 5531 Å^2^ wide seastar-shaped cavity (Fig. 2*B* and *SI Appendix*, Fig. S7*A*).

In all oligomers, the contacts between A_2_B_2_ tetramers are always mediated by B-subunits as shown by rigidly fitting the oxidized A_2_B_2_ crystal structure (PDB ID 2PKQ) (21) inside their respective cryoEM density maps (Figs. 2*B*, 3*A, B* and *E*-*H* and *SI Appendix*, Figs. S6*A*-*C* and S7*A*-*C*). Although A- and B-subunits show a high sequence identity (about 81%; *SI Appendix*, Fig. S8) and similar overall structure, the positioning of B-subunit rather than A-subunit at the contact regions between adjacent tetramers, gave significantly higher correlation coefficients (*SI Appendix*, Table S3). Consistently, the capability of AB-GAPDH to aggregate is long known to depend on the CTE, which is exclusive of B-subunits (15, 34).

### Dissecting the assembling of A_2_B_2_-GAPDH tetramers in higher order oligomers: the role of the CTE

The cryoEM density maps of A_4_B_4_ and both conformers of A_8_B_8_ show unassigned densities in proximity of the contact regions between adjacent A_2_B_2_ tetramers (Fig. 3 and *SI Appendix*, Fig. S6*A*-*C*). These densities start from the last residue of the B-subunit of the crystal structure and continue in the catalytic domain of the closest B-subunit of the adjacent tetramer about 20 Å far away. In some cases the density was clearly visible and continuous, in others was less defined. A model of the C_α_ backbone of the CTE, including the side chains of Cys349 and Cys358 forming the regulative disulfide bridge, was built on the basis of the electron density map of the A_8_B_8_-GAPDH main conformer. The model consists of an extended linker region visible in the electron density maps at lower density threshold, followed by a helix, a circular motif determined by the disulfide bond and a final random coil region (Figs. 3*B, F, H* and 4*A*). In all GAPDH oligomers the CTEs mediate the connection between B-subunits belonging to adjacent A_2_B_2_ tetramers, and each tetramer is connected with the adjacent one by two CTEs. The CTE belonging to one B-subunit penetrates into the catalytic domain of the B-subunit of the adjacent tetramer whose CTE in turn enters into the catalytic domain of the B-subunit of the first tetramer in the opposite direction (Figs. 3*A, B* and *D*-*H* and *SI Appendix*, Figs. S6*B* and *C*). The catalytic sites of the A-subunits, two per tetramer, remain free.

**Figure 4.**
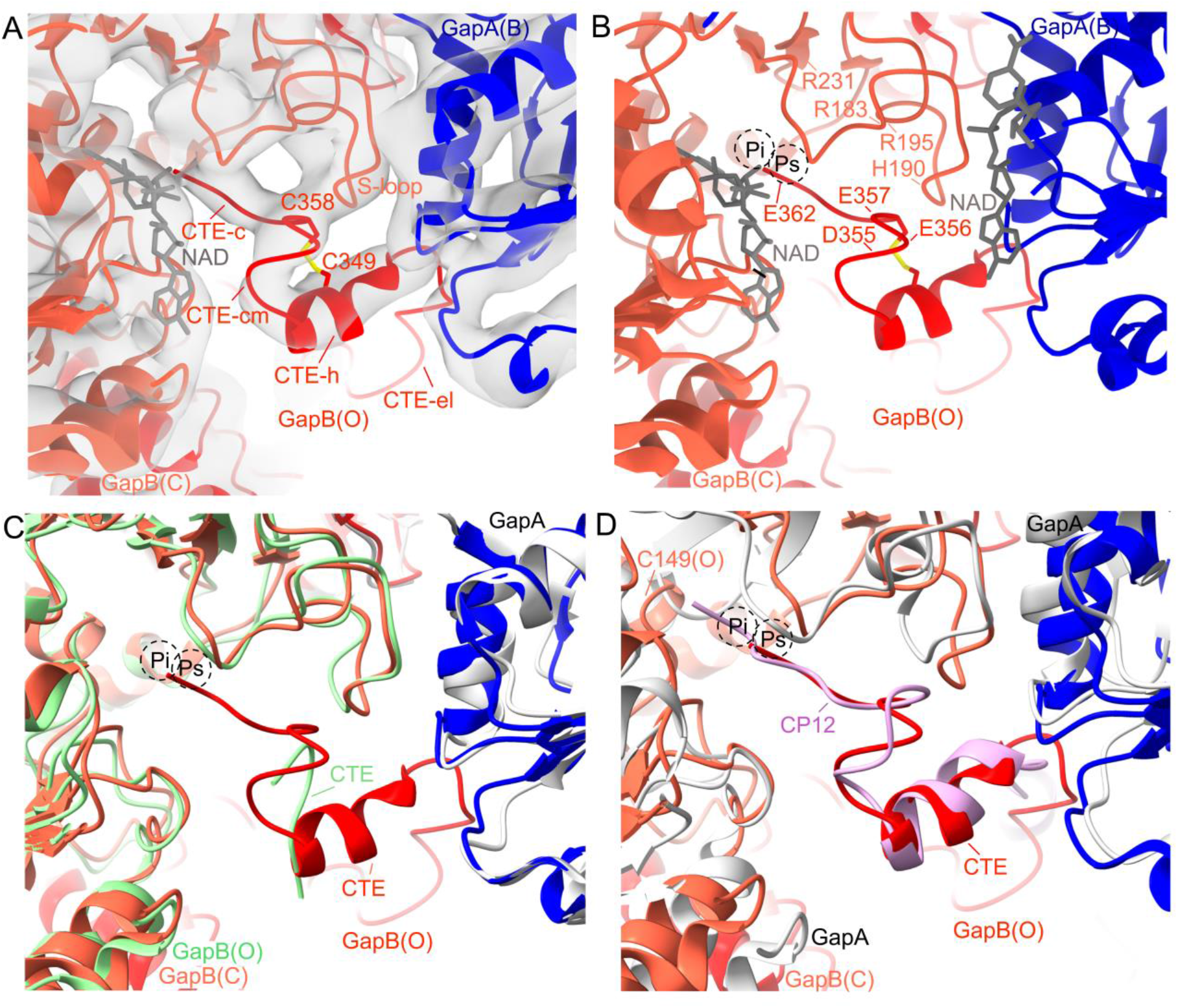
The CTE in the A_8_B_8_ oligomer. (A) Detail of the CTE of B-subunit (chain O) in red inserted in the active site of B-subunit (chain C) in tomato, of the adjacent A_2_B_2_ tetramer. The A-subunit (chain B) is in blue. CTE-el: CTE extended linker; CTE-h: CTE helix; CTE-cm: CTE circular motif; CTE-c: CTE random coil. (B) Detail of the CTE of B-subunit (chain O) in red inserted in the active site of B-subunit (chain C) in tomato, of the adjacent A_2_B_2_ tetramer. The A-subunit (chain B) is in blue. The negatively charged residues of CTE likely interacting with the positively charged residues of B-subunit are indicated. The NAD^+^ bound to the A-subunit is also shown. The P_s_ and P_i_ labels indicate the substrate binding site. (C) Detail of the CTE of B-subunit (chain O) in red superimposed to the CTE of B-subunit (chain O) in green from the crystal structure of the oxidized A_2_B_2_ complexed with NADP^+^ (PDB ID code 2PKQ) (21). The B-subunit (chain O) and the A-subunit of A_2_B_2_ crystal structure are in green and light grey, respectively. Colour code for cryoEM structure is as in panels (A) and (B) Note that the two CTEs shows a different conformation and the CTE from A_2_B_2_ crystal structure ends in the more external region of the catalytic cavity, far away the substrate binding site (P_s_ and P_i_ sites). (D) Detail of the CTE of B-subunit (chain O) in red superimposed to the CP12 C-terminal domain in violet, from the cryoEM model of the ternary GAPDH-CP12-PRK complex (PDB ID 6GVE) (27). The A-subunits of GAPDH from the complex crystal structure are shown in light grey. The catalytic Cys149 is indicated. Note that CTE and the C-ter domain of CP12 have a very similar conformation and CP12 fills both the P_s_ and the P_i_ sites differently from CTE which ends in the P_s_.

The CTE linker regions (Figs. 3*B, F, H* and *SI Appendix*, Fig. S6*C*) differ significantly from each other in length (from 15 Å to 22 Å) and conformation among and inside the different oligomers. They are indeed highly flexible as corroborated by ResMap results that pointed to a significant decrease in resolution in the CTE linker regions (Fig. 3*C, I* and *SI Appendix*, Fig. S6*D*).

The A_2_B_2_ and A_4_B_4_ are able to form higher oligomers, having two “non-engaged” CTEs that are likely free to move in the surroundings and therefore not observed in their corresponding electron density maps (Fig. 3A and *SI Appendix*, Fig. S4*A*). Consistently, the chimeric form composed of A-subunits fused with CTE [(A+CTE)_4_] makes oligomers that reach an unexpectedly high molecular mass, at least 7-fold bigger than the corresponding tetramer (15, 34).

Considering that the A_8_B_8_ oligomer shows each available CTE engaged with another B-subunit (Figs. 3*D*-*H* and *SI Appendix*, Fig. S6*A*-*C*), it is suggested to be the end-point of the oligomerization process. A similar situation is probably present in the A_10_B_10_, but the limited resolution of the electron density map prevented the CTE reconstruction (*SI Appendix*, Fig. S7*A*-*D*).

The last portion of the CTE (helix, circular motif and terminal random coil) of each B-subunit penetrates into the catalytic site of a B-subunit of the adjacent tetramer through the large cavity formed between the bound cofactor NAD(H) and its S-loop (Fig. 4A), ending in the P_s_ site that hosts the phosphate group of the substrate and very close to the hydroxyl groups of the nicotinamide ribose (Fig. 4*B*). Therefore, the CTE prevents the access and binding of the substrate in the B-subunit active site. Arginines 195 and 231 involved in the stabilization of the P_s_ site (21), likely interact with the backbone and side chain carboxylate group of CTE C-terminal Glu362. Additional interactions are likely formed between the positively charged residues of the S-loop such as Arg183 and His190 and negative residues of the CTE (Asp355, Glu356 and Glu357) (Fig. 4*B* and *SI Appendix*, Fig. S8). Moreover, Glu356 and Glu357 are located close to the ribose hydroxyl groups of the NAD adenine moiety of the adjacent A-subunit and their extended and negatively charged side chain can possibly interfere with the correct positioning of the NADP+ 2’-phosphate group. This explains why the enzyme needs to replace NADP(H) by NAD(H) in order to assemble in oligomers and why the phosphate cofactor helps the oligomers dissociation.

The cavity occupied by the CTE in A_8_B_8_ cryoEM structure is the same observed in the crystal structure of oxidized A_2_B_2_ complexed with NADP^+^ (PDB ID 2PKQ) (21). In this last structure, it was possible to build only less than ten C-terminal residues of the two CTEs belonging to the B-subunits of the tetramer. Nevertheless, the superimposition of the two structures shows that the last portion of CTE has a different conformation and in oxidized A_2_B_2_-GAPDH complexed with NADP^+^ ends in the more external region of the catalytic cavity leaving free the P_s_ and the P_i_ sites (Fig. 4*C*) (21, 38).

The CTE responsible of all regulatory properties of A_2_B_2_-GAPDH, is considered evolutionary derived from CP12, being homologous to the C-terminal domain of CP12 (15, 32). The structural models of the binary A_4_-GAPDH/CP12 and ternary A_4_-GAPDH/CP12/PRK complexes (12, 27, 39, 40) reveal that the CTE in A_8_B_8_-GAPDH and the C-terminal domain of CP12 share not only the same cavity but also a very similar conformation (Fig. 4*D*). Indeed, the C-terminal domain of CP12 fits well inside the unassigned density of each B-subunit and particularly the α-helix portion appears well superimposable. The unique striking difference is that CP12 penetrates more deeply in the GAPDH active site compared to CTE, blocking both P_s_ and the P_i_ sites. Indeed, the side chain of Asn78, the last CP12 residue, is observed at an H-bond distance from the thiol group of the catalytic Cys149 (12, 39, 40).

### Interface analysis of AB-oligomers

The A_2_B_2_ tetramers within oligomers are linked together by the CTEs but appear to interact also through a different surface. PDBePISA (41) calculations show that in all GAPDH oligomers the CTEs contribute to the interface area between A_2_B_2_ tetramers by 39% in A_4_B_4_, 32% and 33% in A_8_B_8_ and its alternative conformer, respectively (*SI Appendix*, Table S4).

A_8_B_8_ oligomer shows the largest total interface area (2641 Å^2^) and consequently the largest average single interface area equal to 660 Å^2^ (449 Å^2^ without CTE). This area decreases to 625 Å^2^ (421 Å^2^ without CTE) in the case of the alternative conformer and to 656 Å^2^ (403 Å^2^ without CTE) for A_4_B_4_. The A_10_B_10_ has the smallest average single interface area (228 Å^2^).

CTE-independent interacting surfaces are similar in all oligomers and invariably include four stretches of residues (77-80; 97-114; 119-127; 139-143) located in α-helices and loops (Fig. 3, *SI Appendix* Figs. S6*C* and S8). The last two stretches contain two amino acid insertions in B-compared to A-subunit (Ser123A and Val140) and various sequence differences (*SI Appendix* Fig. S8). This may explain (42) why artificial tetramers made of B-subunits only (B_4_) or (A+CTE)_4_ form oligomers of different size under inactivating conditions (491 *vs* >1800 kDa, respectively) (15, 32, 34).

In A_4_B_4_ and A_8_B_8_ oligomers, but not in A_10_B_10_, additional interface regions comprise residues from the S-loop (179-195) and residues between strands β_2_ and β_3_ (206-208 and 215-222).

Intriguingly, the CTEs also play a key role in improving the thermodynamic stability of both A_4_B_4_ and A_8_B_8_ oligomers. The calculated dissociation free energy (ΔG_diss_) is negative in all oligomers without CTEs indicating that they are unstable, while the presence of CTE prevents their dissociation (*SI Appendix*, Table S4). The most stable oligomer is A_8_B_8_ in the main conformation (ΔG_diss_ = 41 kcal/mol), followed by A_4_B_4_ (ΔG_diss_ = 35.9 kcal/mol) and the hexadecamer alternative conformer (ΔG_diss_ = 35.5 kcal/mol).

### SEC-SAXS data matching with AB-structural models

The theoretical scattering profiles of cryoEM models of the AB-GAPDH oligomers (here presented), and the A_2_B_2_ crystal structure (PDB ID 2PKQ) (21) were calculated (*SI Appendix*, Fig. S9) to evaluate the agreement with SEC-SAXS data and the contribution of the different oligomers.

The inactive sample relative abundance (particles percentage of 19%, 49%, 30% and 2% for A_10_B_10_, A_8_B_8_, A_4_B_4._ and A_2_B_2_,respectively) shows a general agreement with the cryoEM data, except for the negligible contribution of A_2_B_2_ and a larger fraction of A_10_B_10_ (Fig. 2*B*; *SI Appendix*, Fig. S10*A, D*). The comparison between the theoretical and experimental scattering profiles suggests that the data from the inactive sample can be also interpreted reasonably well in terms of one prevailing oligomer at their elution maxima i.e. A_10_B_10_, A_8_B_8_ and A_4_B_4_(*SI Appendix*, Fig. S10*E*; grey *vs*. black line). The A_4_B_4_ coexists with the predominant A_8_B_8_ in a rapid exchanging process and its scattering became dominant only at the tail of the elution (*SI Appendix*, Fig. S10*A*).

Data from the active sample are well interpreted by the scattering profile of the A_2_B_2_ tetramer (*SI Appendix*, Fig. S10*C, D* and *G* and Table S5), while the active-short sample consists of a more complex mixture, predominantly composed by the A_2_B_2_ form coexisting with a significant fraction of A_4_B_4_ oligomer and AB dimers (*SI Appendix*, Fig. S10*B, F*). The introduction of this last form already described for non-photosynthetic GAPDHs (43, 44), clearly improved the fitting (*SI Appendix*, Fig. S10*F;* black *vs*. grey line). However, the absence in the experimental data of the pronounced minimum observed at q=0.1 Å^-1^ in the A_2_B_2_ theoretical scattering profile, can also be ascribed to a quaternary structure rearrangement in solution, that generates a less compact and isometric tetramer (26, 45).

### Concentration effect on the oligomerization of AB-GAPDH

SAXS measurements without SEC separation (SC-SAXS) on AB-GAPDH in inactive and active conditions were also performed (*SI Appendix*, Table S6). The inactive sample can be described as a mixture in which the A_10_B_10_ oligomer is predominant (roughly 50% volume fraction), coexisting with the A_8_B_8_ oligomer (35%) and a smaller fraction of the A_4_B_4_ form (15%) (*SI Appendix*, Fig. S11*A, B* and *C*).

In the active sample, a systematic decrease of the average dimensions and forward scattered intensity was observed with the decrease of the protein concentration (*SI Appendix*, Table S6). The P(r) functions underwent a systematic decrease of the additional peak at 100 Å seen in the bimodal P(r) of higher oligomers, in favor of the main peak at 50 Å characteristic of the A_2_B_2_ tetramer (*SI Appendix*, Fig. S11*D*). The data fitting in terms of a mixture suggests that the fraction of A_2_B_2_ increased from roughly 20% to above 60% upon dilution, at the expenses of the A_4_B_4_ and A_10_B_10_ oligomers, present as 50% and 27% volume fractions, respectively, in the most concentrated sample (*SI Appendix*, Fig. S11E, *F* and Table S7).

This analysis shows that the cryoEM models explain a consistent amount of the SAXS signal. However, the AB-GAPDH oligomerization equilibrium in solution appears more complex. Indeed, partially formed oligomers or less symmetric conformations of (A_2_B_2_)_n_ (n=4 and 5) oligomers such as polymeric chains of A_2_B_2_ units with free CTEs, and small fractions of larger assemblies (n>5), could explain the non-optimal agreement of the fits based on the cryoEM models only and the maximum sizes larger than 240 Å (expected for the A_10_B_10_ oligomer) detected in the inactive sample.

### Concluding remarks

NAD(P)H-dependent GAPDH enzymes are involved in photosynthetic carbon assimilation of all oxygenic phototrophs. However, whereas cyanobacteria and most eukaryotic algae exclusively present a homotetrameric form (A_4_-GAPDH), the major chloroplast GAPDH isozyme of land plants is formed by A and B subunits, the latter containing a redox-sensitive C-terminal extension (CTE) which controls the NADPH-dependent activity of the enzyme and the capability to form higher order oligomers (15, 32).

In this study, we have structurally characterized photosynthetic AB-GAPDH and disclosed the CTE-mediated regulation/oligomerization process, by combining SEC-SAXS and single-particle cryoEM analysis. Both experimental approaches highlighted the presence in both active and inactive *in vitro* conditions (mimicking light and dark *in vivo* conditions) of various oligomers in addition to the expected species with A_2_B_2_ and A_8_B_8_ stoichiometries, respectively (15, 17, 21, 28). In activating conditions beside the heterotetramer A_2_B_2_, the octamer A_4_B_4_ was detected, while in inactivating conditions the population increases to four species, i.e. (A_2_B_2_)_n_ with n=1, 2, 4 and 5 (Figs. 1 and 2). The unexpected dynamism of the AB-GAPDH system is not simply ascribable to the experimental conditions. Indeed, A_4_B_4_ oligomers were observed in leaves of different plant species (35, 46), indicating that this form is both an intermediate step in GAPDH oligomerization and an essential player in its regulation. Moreover, being A_4_B_4_ a structural unit of A_8_B_8_ and likely of A_10_B_10_ oligomers, it represents for the AB-GAPDH system a ubiquitous reservoir of inactive A_2_B_2_ tetramers that when needed can easily dissociate to form the active species or aggregate in higher molecular weight oligomers.

In all oligomers, the interfaces between A_2_B_2_-tetramers uniquely involve B-subunits (Figs. 2*B* and 3), confirming that the CTE manages the AB-GAPDH assembly process upon NADP(H)/NAD(H) cofactor exchange. Moreover, the higher resolution A_4_B_4_ and A_8_B_8_ cryoEM models show that pair of B-subunits from adjacent tetramers hug each other through their CTEs (Figs 3*A, B, E*-*H*, *SI Appendix* Fig. S6*B, C*). Each CTE slips into the cofactor cavity of the partner B-subunit up to its catalytic P_s_ site, effectively preventing the substrate binding (Fig. 4). This positioning of the CTE is only possible if NAD(H) is bound to the A-subunit. However, NAD(H) does not promote oligomerization directly, but it does so by replacing NADP(H). Indeed, the 2’-phosphate of NADP(H) is apparently incompatible with the allocation of the CTE in the active site of B-subunits, justifying the disassembling role of this cofactor (15). On the other hand, the catalytic sites of A-subunits are free and likely available to perform the constitutive NADH-dependent catalysis.

The conformation assumed by the last portion of the CTE closely resembles that one of the CP12 C-terminal domain in the GAPDH-CP12-PRK ternary complex (PDB ID 6GVE) (27) (Fig. 4*C*), indicating that the molecular strategy for the modulation of GAPDH activity appears conserved among all photosynthetic GAPDHs.

In conclusion, our structural study provides a full picture at molecular level showing how the dynamic changes in the oligomeric status of AB-GAPDH contribute to the modulation of the Calvin-Benson cycle in response to light conditions occurring in the natural environment.

## Materials and Methods

### Preparation of AB-GAPDH oligomers

AB-GAPDH isoforms were prepared from partially purified spinach chloroplasts, following ammonium sulfate precipitation, cold acetone precipitation and anion exchange chromatography, as described in (21). Active (NADP^+^-bound) and inactive (NAD^+^-bound) AB-GAPDH samples were obtained as reported in *SI Appendix, Materials and Methods*.

### Small Angle X-ray Scattering data collection and analysis

SAXS data were collected at the BioSAXS beamline BM29 at the European Synchrotron Radiation Facility (ESRF), Grenoble (France) (47). Details of the samples composition, experimental procedures and data acquisition parameters and analysis are reported in *SI Appendix, Materials and Methods*.

### Theoretical scattering profiles calculation from 3D Data

Coordinates from the crystal structure of A_2_B_2_-GAPDH (PDB ID 2PKQ) (21) and from the cryoEM models of AB-GAPDH oligomers (present work) were used to calculate theoretical scattering profiles as detailed in *SI Appendix, Materials and Methods*.

### CryoEM data collection and analysis

Purified inactive AB-GAPDH sample was adsorbed onto a Quantifoil holey TEM grids and plunge-frozen in liquid ethane with a Vitrobot Mark IV cryo-plunger. CryoEM data were collected on a Tecnai F30 Polara cryo electron microscope at 300 keV. AB-GAPDH oligomers were reconstructed in RELION 3.0 (48, 49) and refined in the real-space. Experimental procedures, data analysis and modeling are reported in *SI Appendix, Materials and Methods*.

### Data Availability

The cryoEM maps of AB-GAPDH oligomers and the coordinates of atomic models generated and analyzed in the current study, have been deposited in the Electron Microscopy Data Bank and in the Protein Data Bank, under accession codes: EMD-13824 and PDB ID 7Q53 for A_2_B_2_; EMD-13825 and PDB ID 7Q54 for A_4_B_4_; EMD-13826 and PDB ID 7Q55 for A_8_B_8_ (main conformer); EMD-13827 and PDB ID 7Q56 for A_8_B_8_ (alternative conformer), EMD-13828 and PDB ID 7Q57 for A_10_B_10_.

## Acknowledgements

We deeply thank Prof. Viorel Nicolae Pavel for his essential suggestions on SAXS experiments and data analysis. This work has been supported by Instruct, project number PID 1829 “Unravelling the pathway of regulation of photosynthetic AB-GAPDH by cryo-EM” funded by the Horizon 2020 programme of the European Union. The high-resolution data were collected at the IBS - Institut de Biologie Structurale in Grenoble (France) with assistance from Dr. Guy Schoehn. We thank the European Synchrotron Radiation Facility for allocation of SAXS beam time (BAG Proposals MX1750) and the staff of beamline BM29 for technical support. S.F. and G.F. thanks the Consorzio Interuniversitario di Ricerca in Chimica dei Metalli nei Sistemi Biologici (CIRCMSB).

