## Supplementary Figures and Tables for "Unravelling the regulation pathway of photosynthetic AB-GAPDH"

**This PDF file includes**:
Supplementary text
Figures S1 to S11
Tables S1 to S9
SI references

**Materials and Methods**

**Purification of AB-GAPDH oligomers.** Active and inactive oligomers were obtained incubating overnight at 4°C pure AB-GAPDH enzyme in the presence of 5 mM reduced DTT, 1 mM $NADP^+$ and 1,3-bisphosphoglirate (obtained by incubation of phosphoglycerate kinase, 20 U $ml^{-1}$, with 15 mM 3-phosphoglyceric acid, 10 mM ATP and 5 mM $MgCl_2$) or 5 mM oxidized DTT and 1 mM $NAD^+$, respectively. Following incubation, samples were separately loaded into a Superdex 200 10/300 GL (Cytiva) column, pre-equilibrated in 25 mM K-phosphate, pH 7.4 plus 0.1 mM $NADP^+$, for the active oligomer, or 0.1 mM $NAD^+$, for the inactive oligomers. Measurements of enzyme activity (1) and hydrodynamic radius, and SDS-PAGE were performed on the fractions of the size exclusion chromatography (SEC) before pooling, change the buffer and concentrate the samples. Protein concentration was measured by means of the BCA assay and samples were stored at -80°C before the analyses.

**Dynamic Light Scattering measurements.** The hydrodynamic radius of AB-samples was measured by Dynamic Light Scattering (DLS) employing a Malvern Nano ZS instrument equipped with a 633 nm laser diode. Samples were introduced in disposable polystyrene cuvettes (100 µL) of 1 cm optical path length. The width of DLS hydrodynamic radius distribution is indicated by the polydispersion index. In the case of a monomodal distribution (Gaussian) calculated by means of cumulant analysis, $PdI = (\sigma/Z_{avg})^2$, where σ is the width of the distribution and $Z_{avg}$ is the average radius of the protein population. The reported hydrodynamic radii ($R_h$) have been averaged from the values obtained from five measurements, each one being composed of ten runs of 10 seconds.

**Small Angle X-ray Scattering Data Collection and Analysis**. In SEC-Small Angle X-ray Scattering (SAXS) experiments, the storage buffer (25 mM K-phosphate, pH 7.5) of the active AB-GAPDH sample contained 5 mM reduced DTT, 20 mM $NADP^+$ and 1,3-bisphosphoglirate, whereas for the inactive AB-GAPDH sample the storage buffer contained 0.1 mM $NAD^+$ (Table S8*A*). For SEC elution, 25 mM K-phosphate, pH 7.5 buffers with 0.1 mM $NADP^+$ or 0.1 mM $NAD^+$ were used for the active and inactive AB-GAPDH samples, respectively (Table S8*A*). An additional sample of the active form, named active-short, was obtained from the inactive sample with an incubation time of 2 hours at room temperature in the presence of 5 mM reduced DTT, 20 mM $NADP^+$ and 1,3-bisphosphoglicerate (Table S8*A*).
SEC-SAXS experiments were performed by loading 100-200 µL of samples, onto a Superdex 200 10/300 GL (Cytivia) column connected to the measurement capillary and pre-equilibrated in 25 mM K-phosphate buffer (pH 7.5) plus 0.1 mM $NADP^+$ or $NAD^+$ to analyze active or inactive AB-GAPDH samples, respectively. The SEC separation was run at a flow rate of 0.5 $mL \cdot min^{-1}$. The UV-vis diode array detector of the HPLC system (Shimadzu) recorded the chromatograms at 280 nm before directing the samples to the capillary for SAXS data collection. SAXS frames obtained by 1 s exposure of the capillary, were acquired continuously. Data collection parameters are reported in Table S8*B*.
The automatic pipeline for SEC-SAXS data analysis implemented at BM29 was used to evaluate the quality of the collected data (2) and contributed to the identification of chromatographic regions with constant scattering profiles. Afterward, a classification of the collected frames as buffer or protein frames was performed on the basis of the SAXS intensity trace; the statistical test implemented in CorrMap (3) aided by visual inspection was used to choose the superimposable buffer intensity profiles. The averaging of the buffer scattering data, the subtraction of the averaged buffer intensity from the protein data and an automatic analysis of the subtracted protein profiles were performed with a Matlab script that uses the tools of the ATSAS package (4) to automatically evaluate the scattered intensity extrapolated at zero angle I(0) and the radius of gyration ($R_g$) via the Guinier approximation, and the pair distance distribution function P(r) via the indirect Fourier transform method implemented in GNOM (5). The frame numbers were converted into retention volumes considering the delay between the injection of the sample into the column and the starting time of the SAXS exposure series. Protein frames giving constant $R_g$ values were scaled to the maximum intensity, checked according to the statistical test (3) and then averaged in order to obtain a single representative scattering profile with a better signal to noise ratio.
In SAXS experiments performed with the automatic sample changer (SC), the active sample was stored in a 25 mM K-phosphate, pH 7.9 buffer containing 1 mM $NADP^+$ (Table S8*C*). The 21.2 $mg \cdot mL^{-1}$ stock was diluted with the same buffer just before the SAXS measurements to obtain a concentration series in the range 0.1-2.0 $mg \cdot mL^{-1}$, estimated from the dilution factors. The inactive samples measured as a concentration series in SC mode were directly stored at the final concentration measured by means of BCA assay (0.39-1.89 $mg \cdot mL^{-1}$) or estimated from the dilution factor (0.08-0.2 $mg \cdot mL^{-1}$) in a 25 mM K-phosphate, pH 7.5 buffer containing 1 mM $NAD^+$(Table S8*C*).
SC-SAXS measurements on AB-GAPDH samples in active and inactive conditions were performed by flushing volumes of 50-60 µL and making a set of 10 consecutive exposures during sample flowing in the capillary. The frames were automatically compared to assess radiation damage and then averaged. The

scattering contribution of the capillary filled with buffer was subtracted and the intensity was divided by the protein mass concentration. The absolute intensity scaling using water scattering as a standard (6) and considering a protein specific volume value of 0.735 $cm^3 \cdot g^{-1}$ provided intensities in kDa units. Two repetitions of the measurement procedure for each protein concentration were run and the data were averaged. Sample details and data collection parameters are reported in Table S8*C*, *D*.
Analysis of the scattering profiles was performed with the tools of ATSAS 2.8 ((4)). The I(0) and the $R_g$ were calculated using the Guinier approximation and the indirect Fourier transform method was applied to obtain the P(r) function, with an estimate of the maximum particle dimension ($D_{max}$), in addition to an independent calculation of I(0) and $R_g$. The molecular weight was estimated from (i) the Porod volume ($V_P$) according to the proportionality empirically found for roughly globular proteins (MW ~ $0.625 \cdot V_P$) (7); (ii) the invariant volume-of-correlation length ($V_c$) through a power-law relationship between $V_c$, $R_g$ and MW that has been parametrized (8); and (iii) a method based on an empirical relation to the Porod invariant estimated with a truncated integral (9). In addition, the approach based on Bayesian inference to estimate a most probable value and a confidence interval from all these concentration-independent methods was applied (10).

**Theoretical scattering profiles from 3D Data.** Theoretical scattering profiles were calculated from the crystallographic coordinates of oxidized $A_2B_2$ (PDB ID code 2PKQ) (11) and from the atomic models of AB-GAPDH oligomeric species obtained by the cryoEM analysis (present work), by using CRYSOL 3.0 (4) with default parameters and imposing a q range of 0-0.42 $Å^{-1}$ and 881 data points. The theoretical intensities were scaled to have an I(0) coincident with the squared molecular weight of the simulated constructs and employed for the least-square fitting of experimental SAXS profiles as a linear combination of components in which only the volume fractions are optimized, by means of OLIGOMER (12). The optimized volume fractions were converted into protein mass concentration (c; $g \cdot cm^{-3}$) considering the volume fractions equal to mass fractions $w_i$ (assuming all oligomeric species had the same partial specific volume of 0.735 $cm^3 \cdot g^{-1}$) and by multiplying by the overall protein concentration estimated from the I(0) value in absolute units, according to:

$$c[g \cdot cm^{-3}] = \frac{I(0)[cm^{-1}] \cdot N_A[mol^{-1}]}{\Delta\rho_M^2[cm^2 \cdot g^{-2}] \cdot \sum_i w_i MW_i[g \cdot mol^{-1}]}$$

where $N_A$ is the Avogadro number (6.022 $\cdot 10^{23}$ $mol^{-1}$), $\Delta\rho_M^2$ is the squared scattering contrast per mass of protein ($5.04 \cdot 10^{20}$ $cm^2 \cdot g^{-2}$) and $MW_i$ are the molecular masses of the oligomeric components.
An estimate of the contribution of each oligomer in the overall SEC-SAXS elution was obtained by summing up the optimized concentrations of each oligomer for all frames. In order to compare it to the CryoEM particle statistics, this result was also expressed as particle percentage by dividing each overall mass concentration by the MW of each oligomeric component:

$$\%particle_i = \frac{\frac{\sum_{frames} c_i}{MW_i}}{\sum_i \left(\frac{\sum_{frames} c_i}{MW_i}\right)} \cdot 100$$

Additional fits of selected SAXS data with the theoretical scattering of single structural components were performed using CRYSOL 3.0 (4) in fitting mode (number of spherical harmonics 25, number of fitted data points 51). The fitted q range was selected to 0.01-0.25 $Å^{-1}$ for the SEC-SAXS data and to 0.01-0.30 $Å^{-1}$ for the SC-SAXS data.

**Negative staining EM**. Purified inactive AB-GAPDH oligomers (0.1 $\mu g \cdot mL^{-1}$ AB-GAPDH in 25 mM K-phosphate buffer, pH 7.5 and 1 mM $NAD^+$) were first analyzed by negative staining. Briefly a 5 µL drop of sample was applied to a previously plasma cleaned 400 mesh copper carbon film grids and stained with 1 wt/v % uranyl acetate solution. Data were collected on a JEM-1011 (JEOL) transmission electron microscope (TEM), with thermionic source (W filament) and maximum acceleration voltage 100 kV equipped with Gatan Orius SC1000 CCD camera (4008 x 2672 active pixels).

**CryoEM sample preparation and data collection**. For cryo-EM grid preparation, a 3 µL droplet of purified inactive AB-GAPDH sample (1 $mg \cdot mL^{-1}$ in 25 mM K-phosphate buffer, pH 7.5 and 1 mM $NAD^+$) was plunge frozen in liquid ethane cooled at liquid nitrogen temperature on glow discharged Quantifoil holey TEM grids (Cu, 300 mesh, 1.2/1.3 um) at 100 % humidity and 4.5° C. The grids were blotted with filter paper for 5 s using a Vitrobot Mark IV cryo-plunger (Thermo Fisher Scientific). Grid vitrification optimization was performed

on a Tecnai F20 (Thermo Fisher Scientific) Schottky field emission gun transmission electron microscope, equipped with an automated cryo-box and an Ultrascan 2kx2k CCD detector (Gatan).
Data collection was performed on a Tecnai F30 Polara cryo electron microscope (Thermo Fisher Scientific, USA) equipped with a Schottky field emission gun operated at 300 kV and using Leginon automated acquisition software (Gatan). A total of 2228 movies were recorded on a K2 Summit direct electron detector (Gatan) in super resolution counting mode at a nominal magnification of 31,000X corresponding to a final pixel size of 1.21 Å (further details are listed in Table S9).

**Cryo-EM image processing**. Beam induced motion correction and dose weighting were performed on the collected 2228 movies using MotionCorr2 (13). Contrast transfer function (CTF) correction was performed using CTFFIND4.1 (14). Any movies containing low figure of merit scores, substantial drift, low contrast, thick/crystalline ice were manually excluded from further analysis. The majority of data processing steps were conducted in RELION 3.0 (15, 16). About 1000 representative particles were manually picked from several averaged micrographs. The obtained low pass filtered 2D class averages have then been used for automated particle picking on a total of 1988 averaged micrographs. This resulted in 253954 particles which were extracted and down-sampled (64 X 64) for several iterative rounds of 2D classification and selection. A total of 127963 particles from 2D classes that possessed the quaternary features of the different GAPDH oligomers were subjected to unsupervised 3D classifications (number of classes K = 8) using two unbiased low resolution initial models (an ellipsoid and a sphere). Each 3D classification resulted in eight 3D classes of which two had the quaternary structures corresponding to $A_{10}B_{10}$ and $A_8B_8$ (classes 7 and 8, Fig. S3*A* top) and to $A_4B_4$ and $A_8B_8$ (classes 3 and 6, Fig. S3*A* bottom), respectively. New analyses were then run separately for each oligomer, including the dissociated $A_2B_2$. This species, although not resulting in the first overall 3D classification, was clearly observed in negative staining and cryoEM micrographs and in the overall 2D classification (Fig. 2*A* and Figs. S2*A*, *B*). For each oligomer an automated particle picking round was repeated with Gautomatch (https://www.mrc-lmb.cam.ac.uk/kzhang/) using as template the low pass filtered 2D projections derived from the corresponding cryoEM electron density maps obtained in the previous 3D classification. After several rounds of 2D classification and selection, a total of 48558, 31023, 64130 and 33067 particles for $A_2B_2$, $A_4B_4$, $A_8B_8$ and $A_{10}B_{10}$, respectively were subjected to a new 3D classification using as initial models their correspondent low pass filtered (40 Å) previously obtained cryoEM electron density maps (Fig. S3*B*). The initial model for the dissociated $A_2B_2$ tetramers was instead calculated from its assigned 2D averages using the initial model generation tool within RELION3.0 (15, 16). After 3D classifications 19636 particles were assigned to the dissociated $A_2B_2$ (K=4), 20777 particles were assigned to $A_4B_4$ (K=4), 34379 to $A_8B_8$ (23611 particles to the main form and 10768 particles to its alternative conformer, K=8) and finally 7352 particles were assigned to $A_{10}B_{10}$ (K=4). These subsets of particles, after being re-extracted at full resolution, were used for the final refinement. We obtained symmetry-constrained maps at 6.7 Å (D2 point group symmetry), 8.9 Å (C1 point group symmetry), 5.7 Å (C2 point group symmetry), 7.1 Å (C2 point group symmetry) and 13 Å (C5 point group symmetry) for $A_2B_2$, $A_4B_4$,, -$A_8B_8$ (both main and alternative conformer) and $A_{10}B_{10}$ oligomers, respectively. Identical maps were obtained for $A_4B_4$, and both $A_8B_8$ conformers, by removing symmetry constraints (i.e. imposing the C1 symmetry) during the refinement with RELION 3.0 (15, 16). The resolution of the final maps was estimated by the 0.143 FSC criterion after a postprocessing procedure. Estimation of the local resolution was done in ResMap (17). Handedness of the reconstructions was determined by fitting the GAPDH oligomeric models (see below) into the obtained maps using the 'fit in map' tool in Chimera 1.15 (18).

**Modelling and Bioinformatics Tools.** The GAPDH oligomeric models were obtained by rigid-body fitting the crystallographic oxidized $A_2B_2$ model (PDB ID 2PKQ) (11) in their corresponding final cryo-electron density maps using the 'fit in map' tool in Chimera (18). The CTEs of the B-subunits belonging to the more resolved GAPDH oligomers cryoEM density maps (i.e. the $A_4B_4$ and $A_8B_8$) were built as $C_\alpha$ backbones using COOT (19). Afterward, the obtained GAPDH models were independently refined into their corresponding cryoEM density maps using iterative cycles of Phenix real space refinement (20) and COOT (19) manual adjustment. Cross correlation analyses, measures of distances, areas and angles, 3D visualizations and rendering were performed using Chimera and ChimeraX (21). GAPDH oligomers protein interfaces, contacts and free energy of assembly dissociation were calculated using PDBePISA (22) and visualized using Chimera (18).

**Data Availability**
The cryoEM maps of AB-GAPDH oligomers and the coordinates of atomic models generated and analyzed in the current study, have been deposited in the Electron Microscopy Data Bank and in the Protein Data Bank, under accession codes: EMD-13824 and PDB ID 7Q53 for $A_2B_2$; EMD-13825 and PDB ID 7Q54 for

$A_4B_4$; EMD-13826 and PDB ID 7Q55 for $A_8B_8$ (main conformer); EMD-13827 and PDB ID 7Q56 for $A_8B_8$ (alternative conformer); EMD-13828 and PDB ID 7Q57 for $A_{10}B_{10}$.

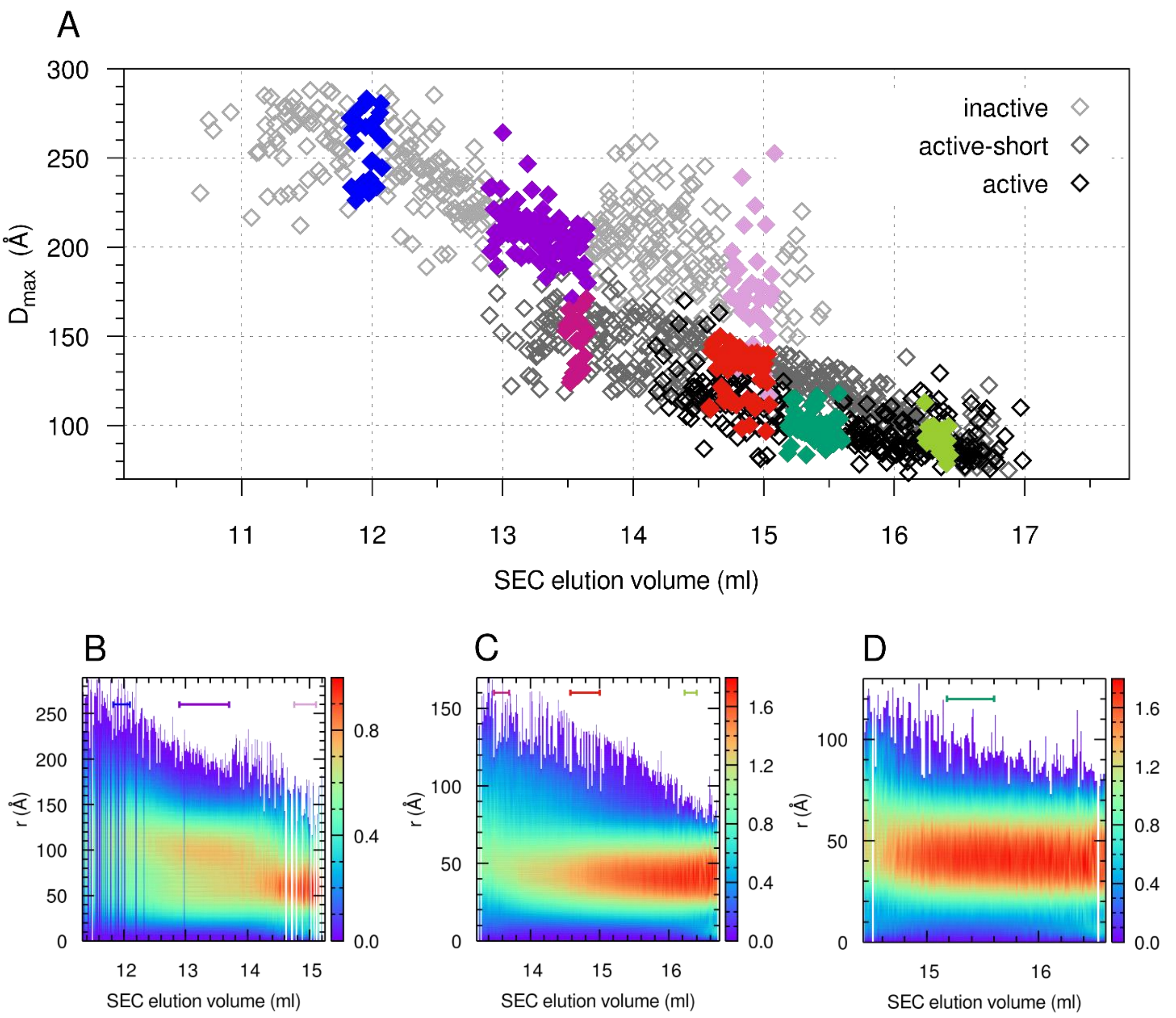


**Fig. S1. SEC-SAXS elution profiles: maximum size and 2D maps for P(r) functions.**
(A) The maximum particle dimension ($D_{max}$, diamonds) estimated from indirect Fourier transform of the SAXS frames for the three AB-GAPDH samples: inactive (light grey symbols, maximum at 13 ml), active-short (grey symbols, maximum at 14.8 ml) and active (black symbols, maximum at 15.4 ml), is shown as a function of the SEC elution volume. The data points belonging to the frames averaged to obtain the selected scattering profiles are highlighted with a colour code.
2D maps of (B) inactive, (C) active-short, (D) active samples analysed by means of SEC-SAXS showing the calculated pair distance distribution function normalized by the subtended area (P(r)/(I(0)), as a function of the SEC elution volume, are presented. The frames averaged to obtain the representative scattering profiles are highlighted by means of bars whose colour key corresponds to that of the plotted P(r) functions in Figure 1.

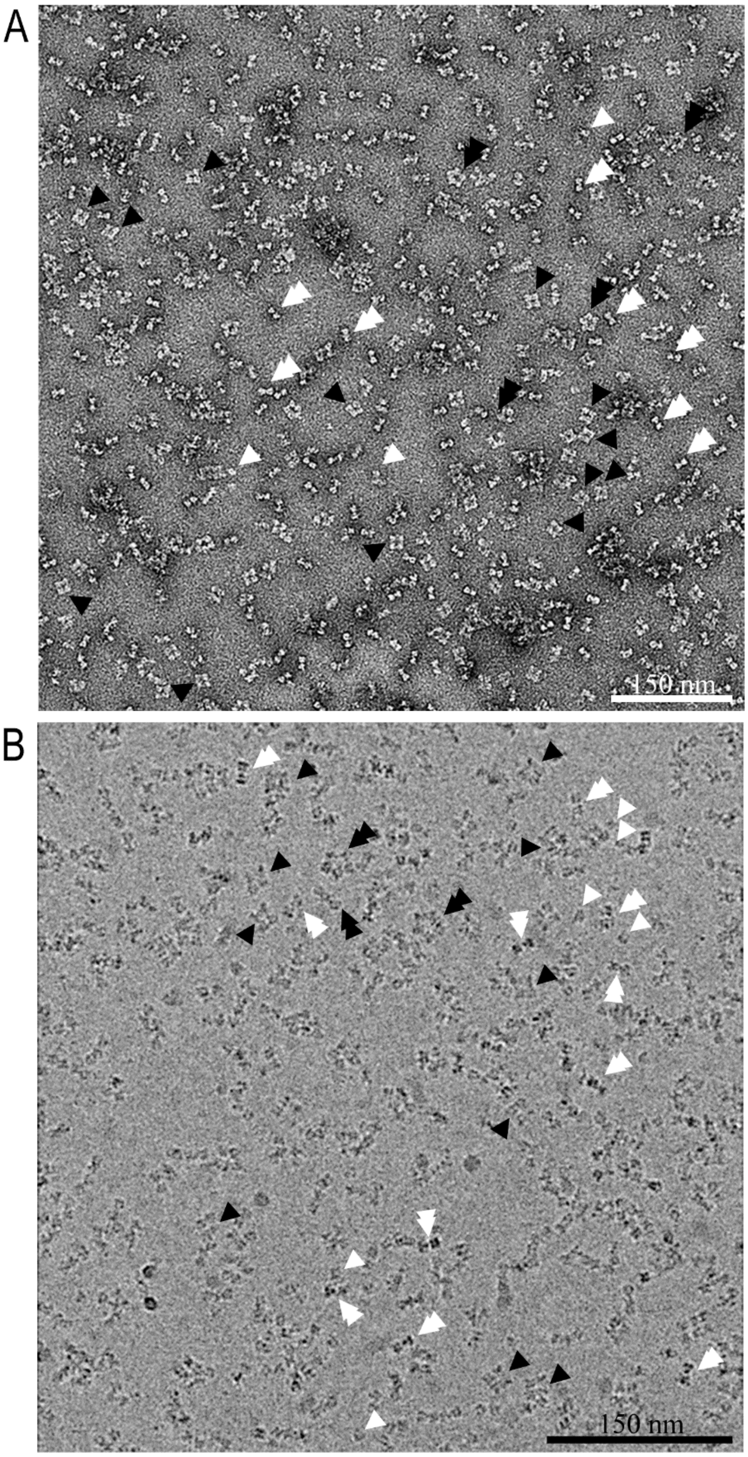


**Fig. S2. Electron microscopy micrographs of GAPDH oligomers in the inactivation buffer.**
(A) Negative staining and (B) cryoEM representative micrographs. The single and double arrowheads point to the $A_2B_2$ (single white arrowheads), $A_4B_4$ (double white arrowheads), $A_8B_8$ (single black arrowheads) and $A_{10}B_{10}$ (double black arrowheads) projections.

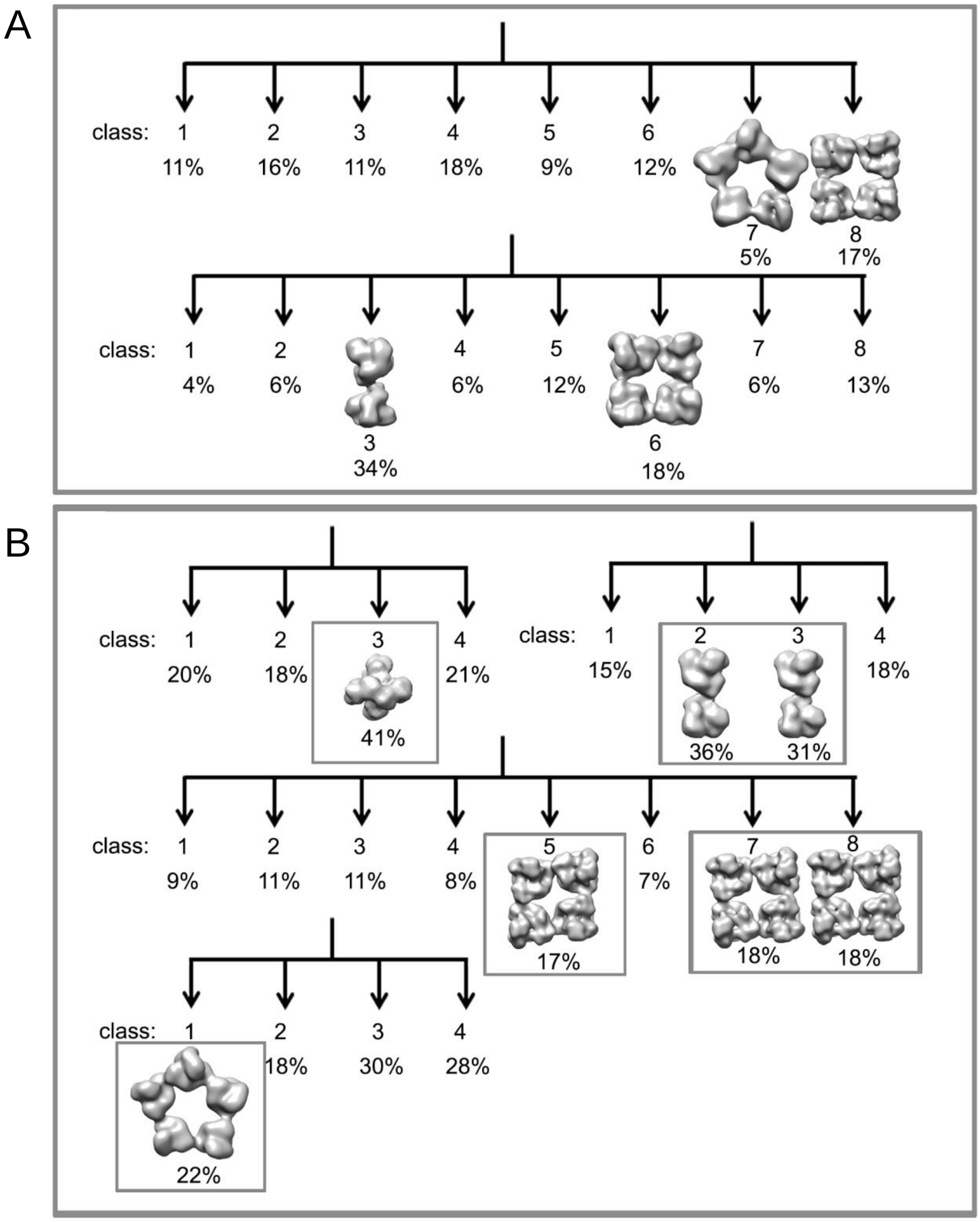


**Fig. S3. 3D classification for GAPDH data set.**
(A) Preliminary 3D classifications performed on the whole GAPDH data set using an ellipsoid (top) and a sphere (bottom) as initial models. (B) 3D classification performed on single GAPDH oligomer data sets. The particles belonging to the boxed 3D classes were used for the final 3D refinement.

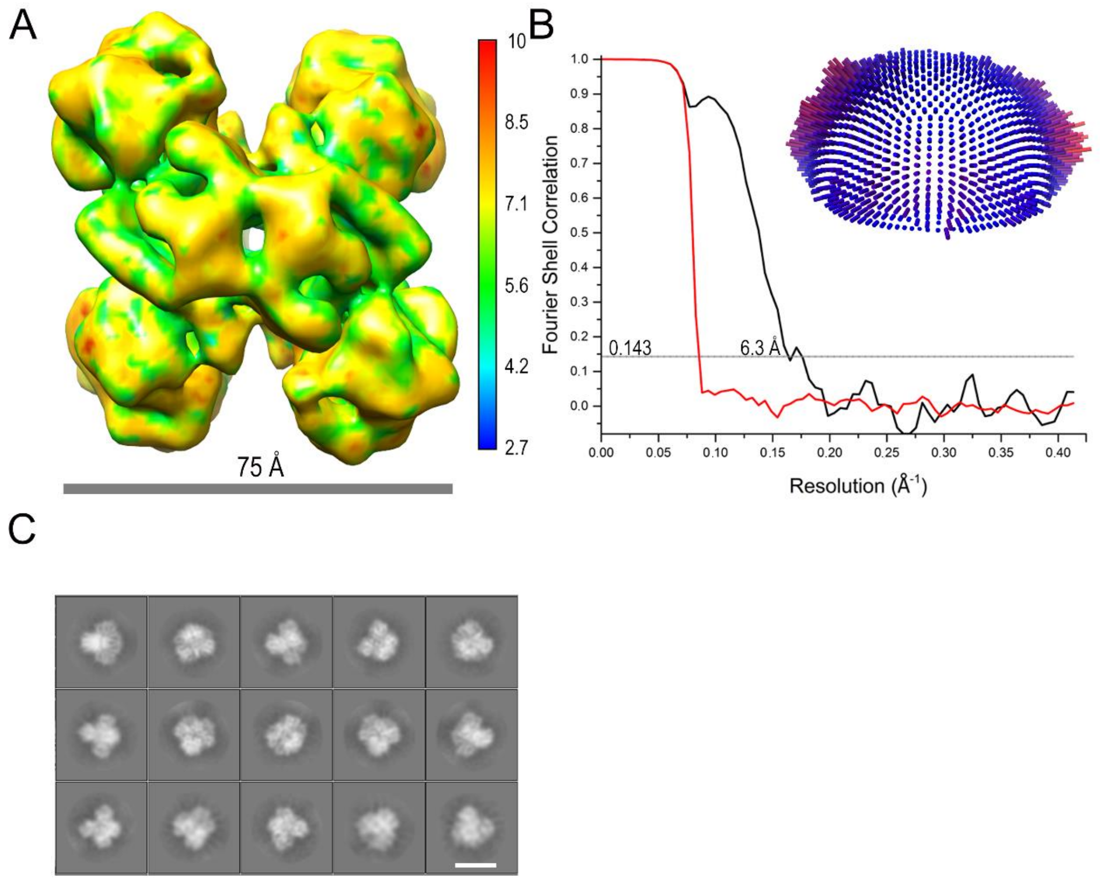


**Fig. S4. The $A_2B_2$ tetramer.**

(A) CryoEM electron density map of $A_2B_2$ oligomer (D2 symmetry) at 6.3 Å resolution filtered according to ResMap local resolution. (B) Fourier shell correlation (FSC) curves (red, FSC phase randomized masked curve; black, FSC corrected curve) of the map with the resolution that corresponds to FSC=0.143 marked. The inset shows the Euler angle distribution. (C) Representative 2D class averages of the $A_2B_2$ particle images. The scale bar is 80 Å.

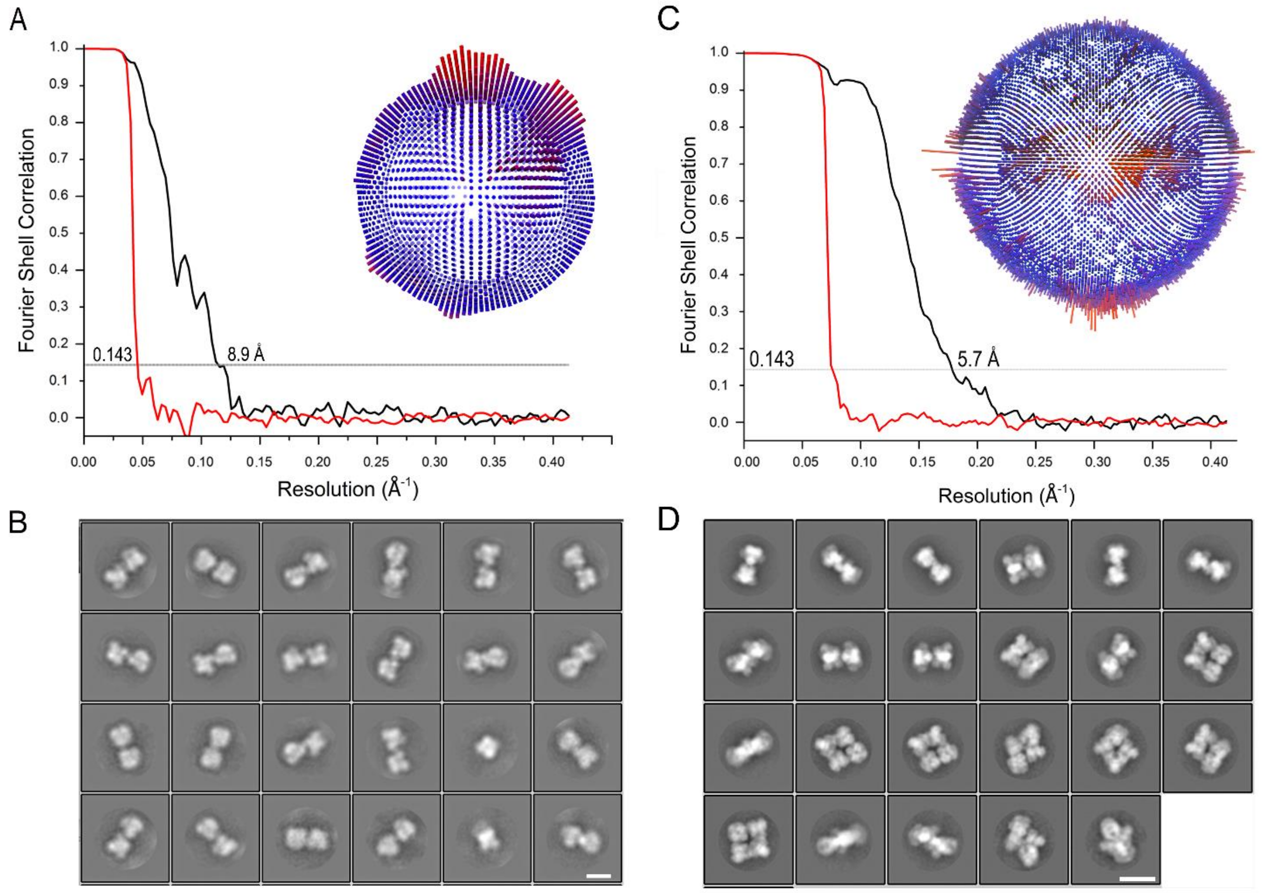


**Fig. S5. FSC curve and 2D classes for $A_4B_4$ and $A_8B_8$ oligomers.**
(A) FSC curve of $A_4B_4$ map (red, FSC phase randomized masked curve; black, FSC corrected curve) with the resolution that corresponds to FSC=0.143 marked. The inset shows the Euler angle distribution. (B) Representative 2D class averages of $A_4B_4$ particle images. The scale bar is 85 Å. (C) FSC curve of the $A_8B_8$ map (red, FSC phase randomized masked curve; black, FSC corrected curve) with the resolution that corresponds to FSC=0.143 marked. The inset shows the Euler angle distribution. (D) Representative 2D class averages of the $A_8B_8$ particle images. The scale bar is 150 Å.

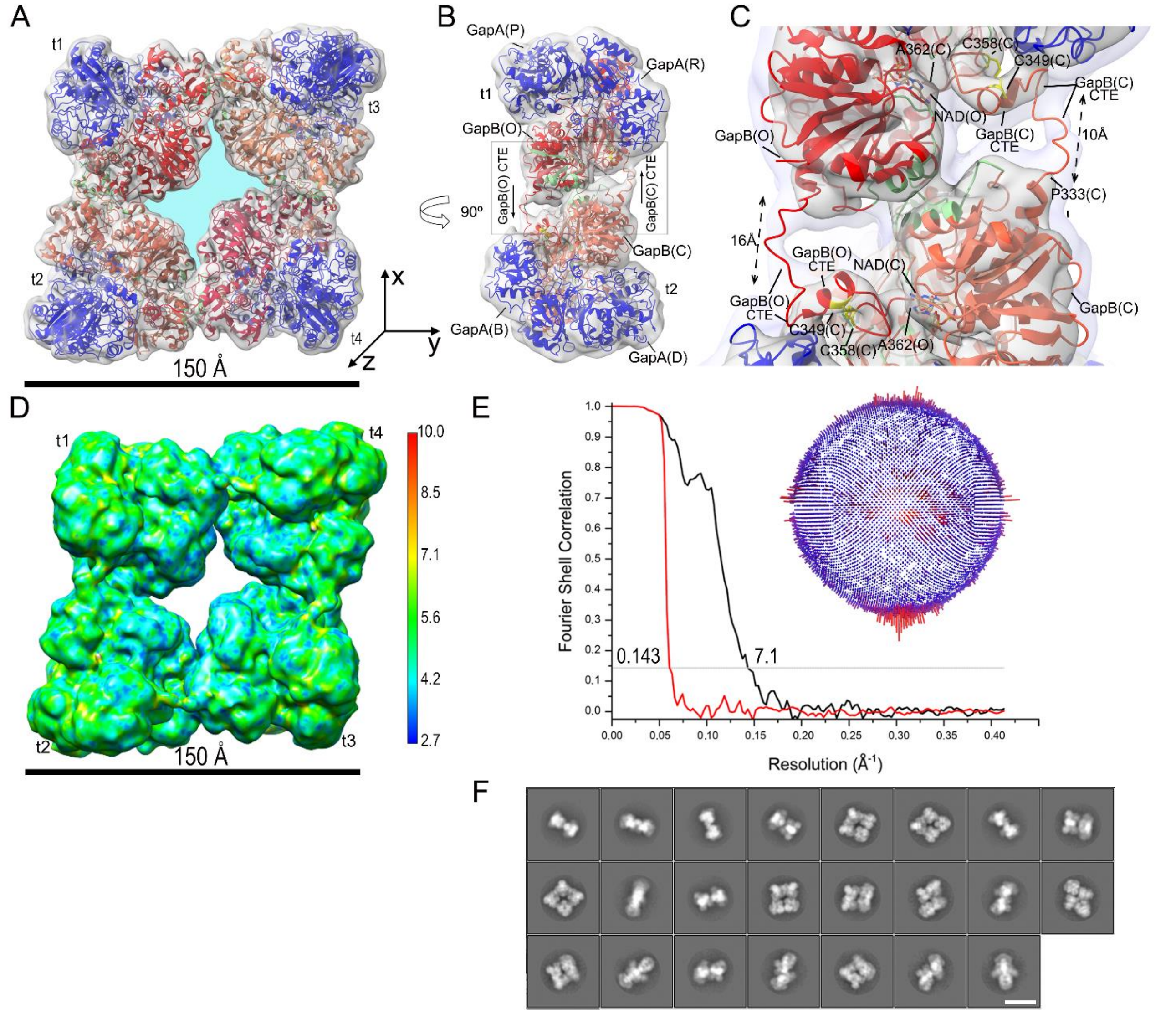


**Fig. S6. The $A_8B_8$ alternative conformer.**
(A) CryoEM electron density map (C2 symmetry) at 7.1 Å fitted with the models derived from the crystal structure of the oxidized $A_2B_2$ complexed with $NADP^+$ (PDB ID 2PKQ) (11). Labels t1-t4 indicate the $A_2B_2$ tetramers. The O/Q, A/C, E/G and K/I B-subunits are represented in red, tomato, crimson and coral, respectively. The A-subunits are in blue. The oligomer central cavity (in light blue) has a surface area of 1738 Å$^2$. (B) Side view of the map in (A) shown at low density threshold. (C) Detail of the region boxed in B. The cryoEM electron density map is displayed at two different isosurface levels (high in dark gray and low in light gray). The interfacing residues between adjacent t1 and t2 GAPDH tetramers are highlighted in green. (D) CryoEM electron density map filtered according to ResMap local resolution. (E) FSC curve of the oligomer map (red, FSC phase randomized masked curve; black, FSC corrected curve) with the resolution that corresponds to FSC=0.143 marked. The inset shows the Euler angle distribution. (F) Representative 2D class averages of the $A_8B_8$ particle images. The scale bar is 150 Å.

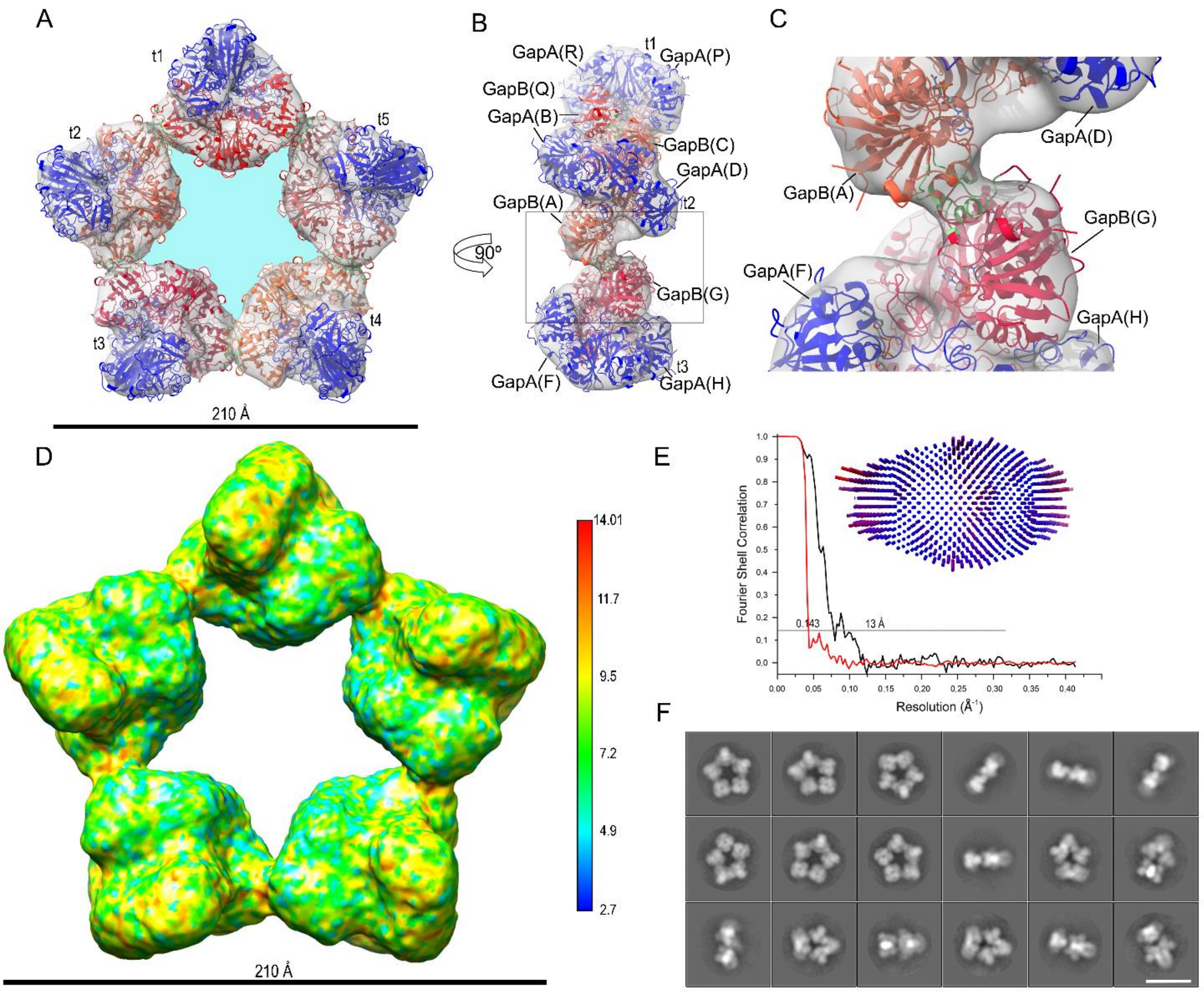


**Fig. S7. The $A_{10}B_{10}$ icosamer.**

(A) CryoEM electron density map (C5 symmetry) at 13 Å fitted with the models derived from the crystal structure of the oxidized $A_2B_2$ complexed with $NADP^+$ (PDB ID 2PKQ) (11). Labels t1-t5 indicate the $A_2B_2$ tetramers. B-subunits are represented in red, tomato, crimson, coral and indian red, while A-subunits are in blue. The oligomer central cavity (in light blue) has a surface area of $5100^2$. (B) Side view of the map shown in A containing the GAPDH tetramers t1-t3. (C) Detail of the region boxed in B. The interfacing residues between B subunits, i.e. B-subunits (chain A) (tomato) and B-subunits (chain G) (crimson) of adjacent t2 and t3 GAPDH tetramers are highlighted in green. (D) CryoEM electron density map filtered according to ResMap local resolution. (E) FSC curve of the oligomer map (red, FSC phase randomized masked curve; black, FSC corrected curve) with the resolution that corresponds to FSC=0.143 marked. The inset shows the Euler angle distribution. (F) Representative 2D class averages of the $A_{10}B_{10}$ particle images. The scale bar is 200 Å.

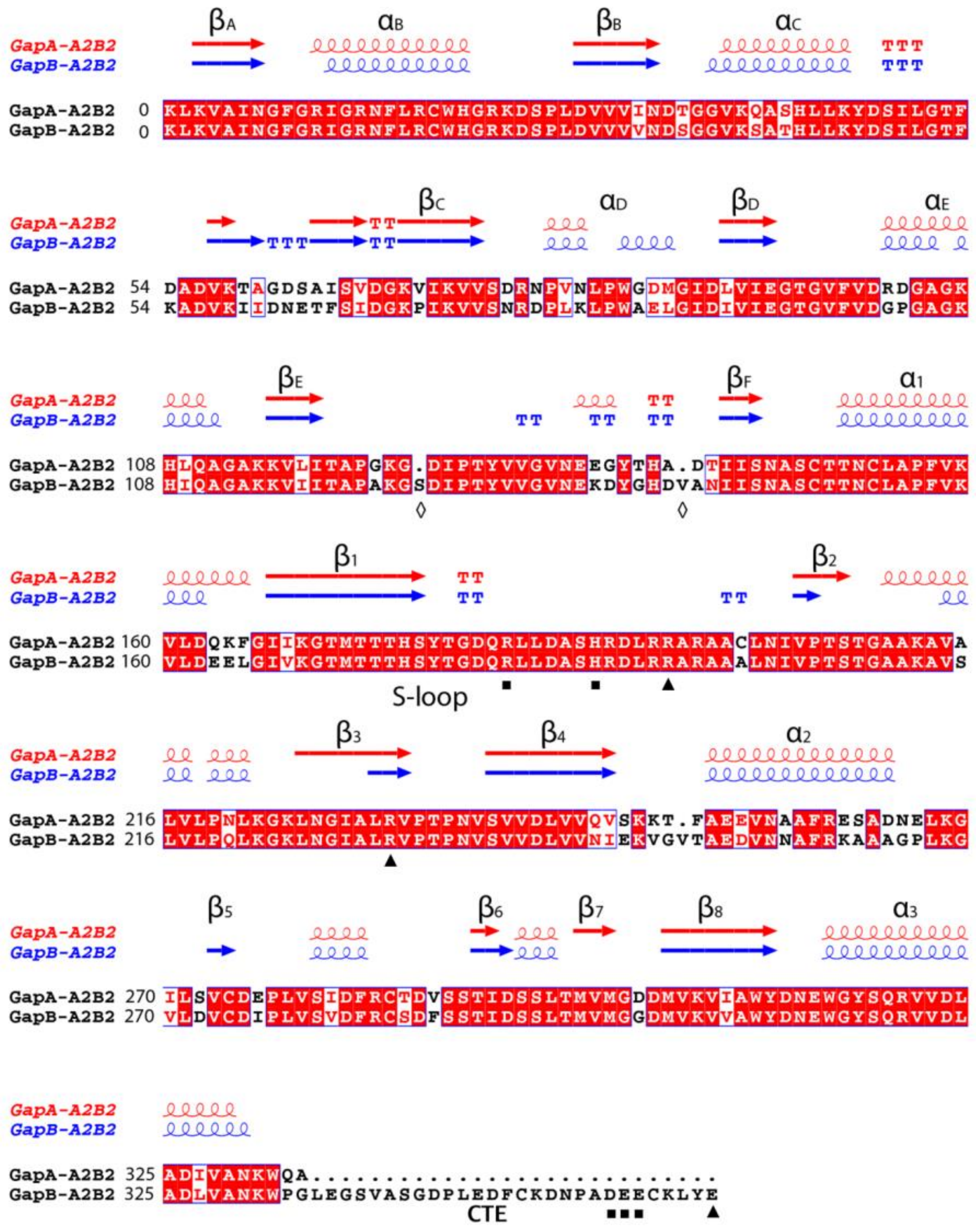

**Fig. S8. Sequence and structural alignment of the A- and B-subunit.**
The alignment was performed with ClustalW and visualized with Esprípt (http://espript.ibcp.fr) using the sequence and the structure of oxidized $A_2B_2$ B-subunit (chain Q) and A-subunit (chain R) (PDB ID 2PKQ) (11). The black squares and triangles indicate residues likely interacting with CTE residues indicated with the same symbols, of the B subunit belonging to an adjacent tetramer (see main text). White diamonds indicate residue insertions of B-subunit respect to A-subunit.

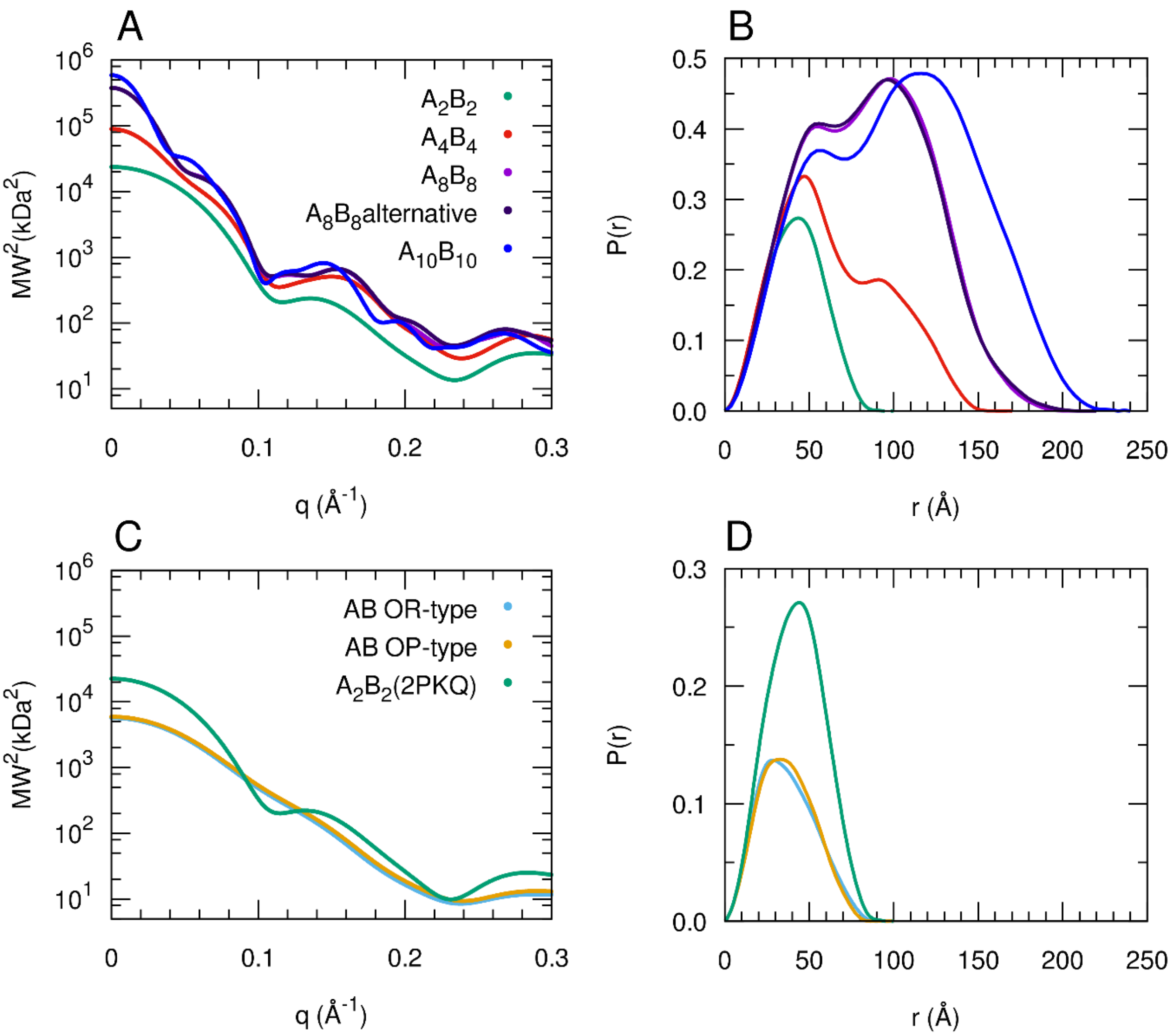


**Fig. S9. Theoretical scattering profiles and P(r) functions of AB-GAPDH oligomers cryoEM models.** (A) Theoretical scattering profiles calculated from the atomic models of the AB-GAPDH oligomers obtained by cryoEM analysis. In (B) the corresponding pair distance distribution functions (P(r)) provided by indirect Fourier transform of the theoretical profiles are shown. (C) Theoretical scattering profiles calculated from the crystal structure of the AB-GAPDH tetramer in oxidized form complexed with NADP (PDB ID 2PKQ) (11) and from two possible dimeric AB forms. In (D) the corresponding P(r) functions are shown.

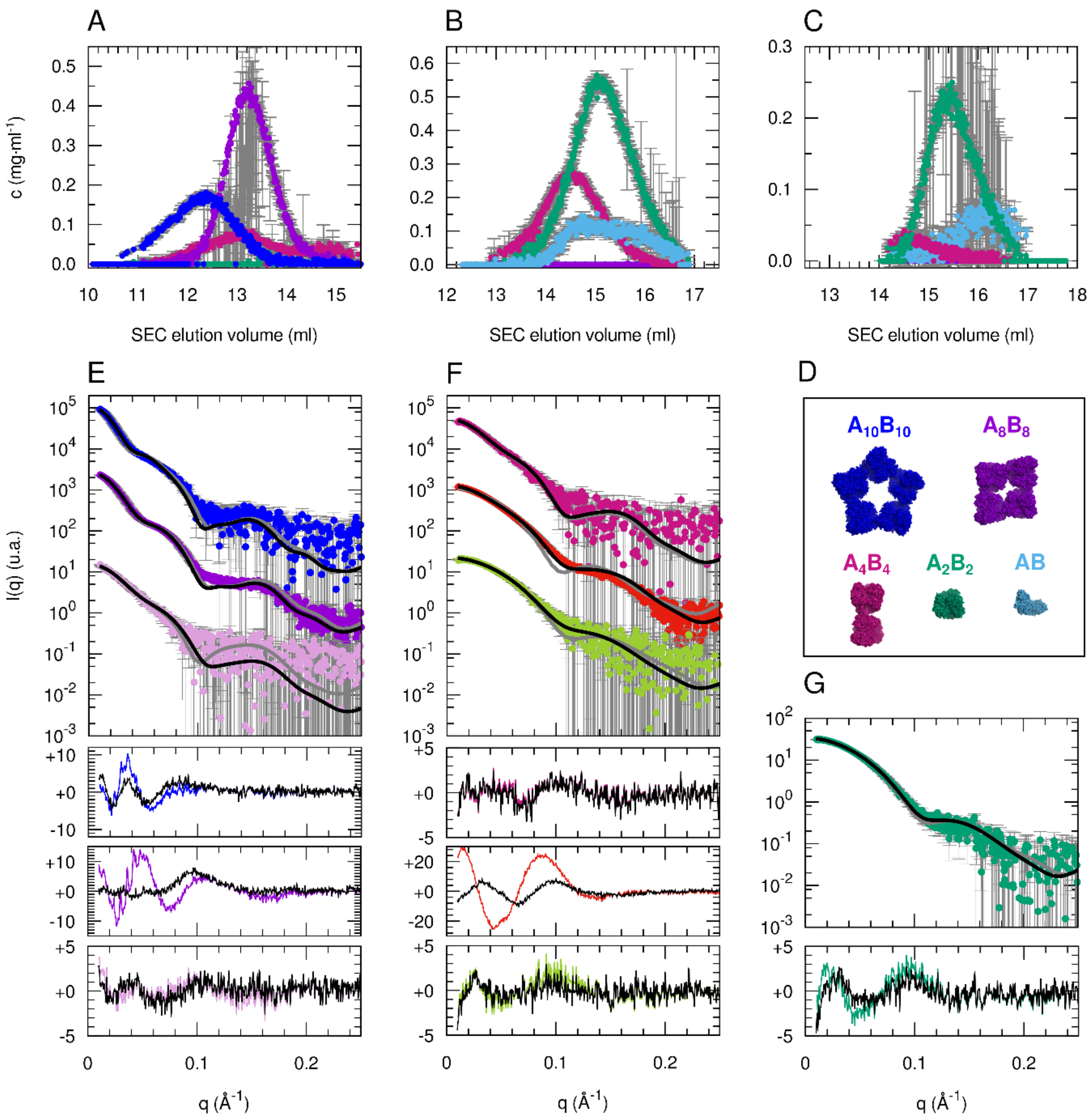


**Fig. S10. Modelling of the SEC-SAXS data on the basis of AB-GAPDH oligomers cryoEM models.** Optimized mass concentrations of the different oligomers as a function of the elution volume (A) for inactive, (B) for the active-short and (C) for active sample. (D) Color code explaination. (E) Best fit of the three selected average SAXS profiles in the elution of the inactive sample (blue, violet and pink circles, colour code as in Figure 1A, B) as linear combinations of the AB-GAPDH oligomers $A_{10}B_{10}$, $A_8B_8$ and $A_4B_4$ (black lines). The optimized volume fractions are reported in Table S5. The best-fits provided by a single atomic structure ($A_{10}B_{10}$, $A_8B_8$ and $A_4B_4$, respectively) are reported as grey lines for comparison. In the panels below error-weighted residual difference plots are reported [$(I_{exp}-I_{calc})/\sigma_{exp}$, where $I_{exp}$ and $I_{calc}$ are the experimental and calculated intensity respectively and $\sigma_{exp}$ are the experimental standard deviations], as black lines for the linear combination fits and as colored lines for the single structure fits. (F) Best fit of the three selected average SAXS profiles in the elution of the active-short sample (purple, red and light green circles, colour code as in Figure 1A, C) as a linear combination of $A_4B_4$, $A_2B_2$ or AB (black lines). The best-fits provided by a single atomic structure ($A_4B_4$, $A_2B_2$ and again $A_2B_2$ respectively) are reported as grey lines for comparison. (G) Best fit of the selected average SAXS profile in the elution of the active sample (green circles, colour code as in Figure 1A, D) as a linear combination of $A_4B_4$, $A_2B_2$ or AB (black line). The best-fit provided by a single atomic structure ($A_2B_2$) is reported as a grey line.

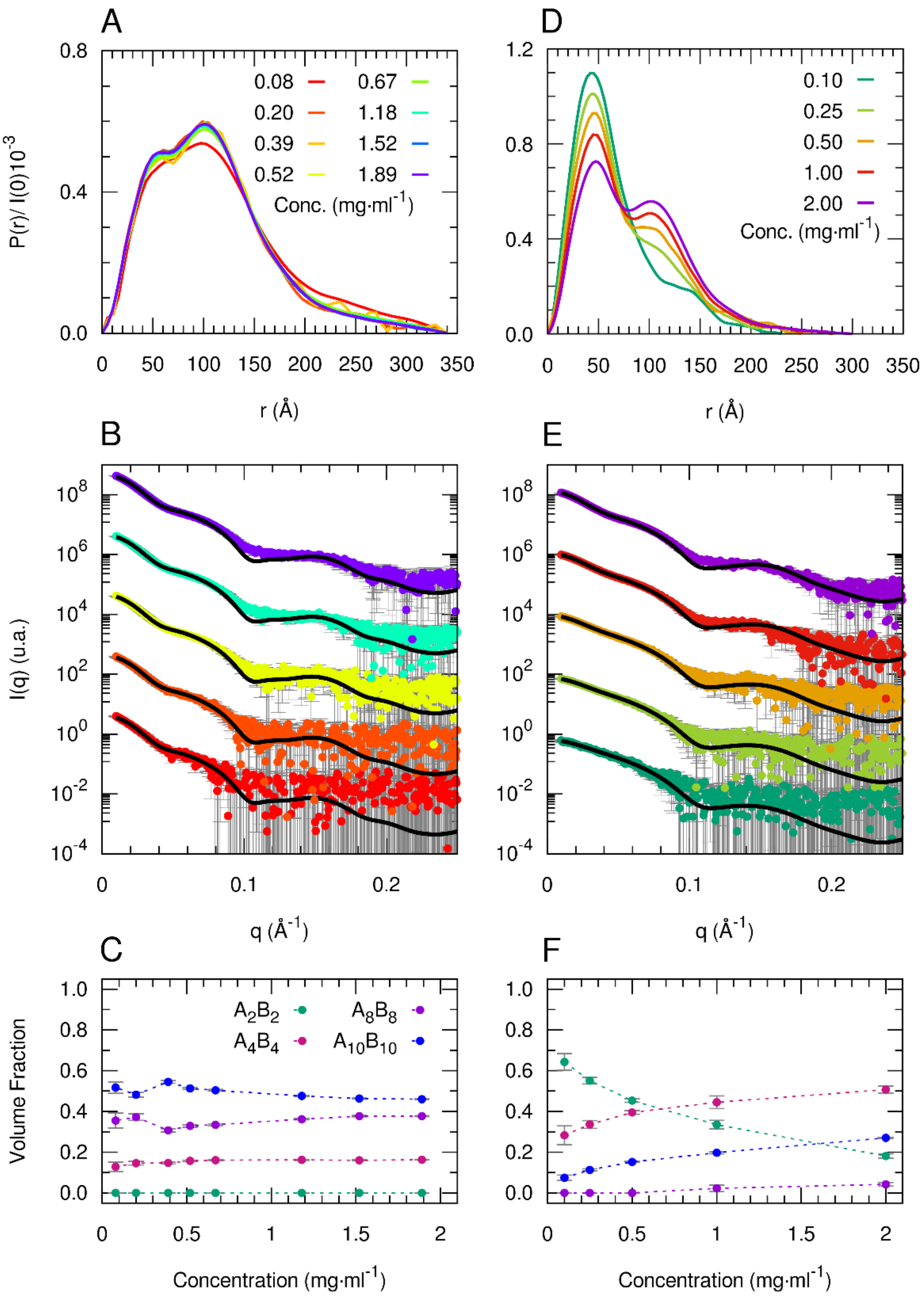


**Fig. S11. SC-SAXS analysis for concentration series of active and inactive AB-GAPDH samples.**
(A) Pair distance distribution functions obtained by indirect Fourier inversion of SAXS data collected on a concentration series of AB-GAPDH incubated in "inactive" conditions. (B) SAXS profiles of a concentration series of AB-GAPDH incubated in "inactive" conditions (dots with colour code reported in (A)) and theoretical scattering profiles (black lines) obtained by fitting the data as a linear combination of the form factors calculated from the atomic coordinates of the $A_2B_2$, $A_4B_4$, $A_8B_8$ and $A_{10}B_{10}$ models presented in the manuscript. (C) Volume fractions of the different AB-GAPDH oligomers as a function of protein concentration, obtained from the fitting of SAXS data reported in (B). In panels (D), (E) and (F) the results of the same SAXS data analysis of AB-GAPDH incubated in "active" conditions are reported.

**Table S1**. Detailed summary of the SEC-SAXS data analysis of AB-GAPDH samples.

| | inactive | | | active-short | | | active |
|---|---|---|---|---|---|---|---|
| SAXS frame at injection | 90 | | | 222 | | | 268 |
| Background data | | | | | | | |
| frames | 1-1295 | | | 1-1650 | | | 1-1886 |
| max $V_{SEC}$ (ml) | 10 | | | 11.9 | | | 13.5 |
| Selected protein data | | | | | | | |
| frames | 1510-1540 | 1638-1728 | 1860-1900 | 1840-1860 | 1972-2027 | 2170-2193 | 2090-2140 |
| $V_{SEC}$ interval (ml) | 11.8-12.1 | 12.9-13.7 | 14.8-15.1 | 13.5-13.7 | 14.6-15.0 | 16.2-16-4 | 15.2-15.6 |
| $\langle V_{SEC}\rangle$ (ml) | 12.0 | 13.3 | 14.9 | 13.6 | 14.8 | 16.3 | 15.4 |
| Guinier fit | | | | | | | |
| $R_g$ (Å) | 80.6 | 66.8 | 59.6 | 50.9 | 39.4 | 34.0 | 33.9 |
| $\sigma(R_g)$ (Å) | 1.0 | 0.1 | 1.5 | 0.6 | 0.1 | 0.4 | 0.1 |
| I(0) [kDa·c(mg/ml)] | 117.9 | 248.5 | 16.0 | 27.2 | 125.8 | 23.3 | 34.0 |
| σ(I(0)) | 0.5 | 0.2 | 0.3 | 0.3 | 0.1 | 0.2 | 0.1 |
| First q point ($Å^{-1}$) | 0.007 | 0.007 | 0.013 | 0.130 | 0.120 | 0.220 | 0.016 |
| Last q point ($Å^{-1}$) | 0.016 | 0.019 | 0.022 | 0.025 | 0.029 | 0.038 | 0.038 |
| Auto$R_g$ quality | 0.89 | 0.87 | 0.96 | 0.74 | 0.97 | 0.99 | 0.96 |
| Indirect Fourier transform | | | | | | | |
| $R_g$ (Å) | 82.7 | 66.9 | 67.2 | 53.2 | 40.3 | 33.9 | 34.1 |
| $\sigma(R_g)$ (Å) | 0.4 | 0.1 | 3.3 | 0.6 | 0.1 | 0.3 | 0.1 |
| I(0) [kDa·c(mg/ml)] | 119.3 | 248.7 | 16.7 | 27.6 | 126.4 | 23.2 | 34.2 |
| σ(I(0)) | 0.4 | 0.2 | 0.5 | 0.2 | 0.1 | 0.1 | 0.1 |
| $V_P$ ($\cdot 10^{-3}$ $Å^3$) | 1100 | 830 | 508 | 346 | 228 | 183 | 201 |
| First q point ($Å^{-1}$) | 0.007 | 0.007 | 0.013 | 0.013 | 0.012 | 0.022 | 0.016 |
| Last q point ($Å^{-1}$) | 0.25 | 0.25 | 0.25 | 0.25 | 0.25 | 0.25 | 0.25 |
| $D_{max}$ imposed for P(r) (Å) | 276 | 217 | 280 | 180 | 146 | 112 | 112 |
| $D_{max}$ variability estimate (Å) | 15 | 10 | 25 | 10 | 10 | 5 | 5 |
| GNOM quality estimate | 0.74 | 0.74 | 0.65 | 0.73 | 0.63 | 0.75 | 0.75 |
| MW($V_P$)[a] (kDa) | 647 | 488 | 299 | 203 | 134 | 107 | 118 |
| MW ($V_c$)[b] (kDa) | 712 | 555 | 222 | 208 | 147 | 112 | 132 |
| MW (MoW)[c] (kDa) | 681 | 568 | 231 | 225 | 173 | 122 | 148 |
| MW Bayesian[d] | | | | | | | |
| estimate (kDa) | 715 | 479 | 318 | 243 | 147 | 119 | 147 |
| estimate probability (%) | 94.6 | 95.0 | 79.1 | 89.0 | 48.8 | 46.5 | 50.4 |
| credibility interval (kDa) | 614-751 | 455-556 | 221-373 | 195-264 | 142-177 | 111-127 | 127-151 |
| interval probability (%) | 99.8 | 98.4 | 99.8 | 99.6 | 98.0 | 92.7 | 95.9 |

[a]From the Porod volume VP according to the empirical relation MW~0.625VP ((7); [b]From the volume of correlation $V_c$ ($q_{max}$ for integration 0.25 $Å^{-1}$) (8); [c]From the Porod invariant ($q_{max}$ for integration 0.25 $Å^{-1}$) (9); [d]From the Bayesian inference approach based on concentration-independent methods (10).

**Table S2.** Summary of dimensional parameters obtained by the analysis of selected SAXS profiles collected during the SEC elution of AB-GAPDH samples.

| | | Guinier | P(r) | | Porod volume | MW estimate | | Possible stoichiometry | |
|---|---|---|---|---|---|---|---|---|---|
| Sample | <$V_{SEC}$> (ml) | $R_g$ (Å) | $R_g$ (Å) | $D_{max}$ (Å)[a] | (·$10^{-3}$ Å$^3$) | MW[b] | MW[c] | MW (kDa) | |
| inactive | | | | | | | | | |
| | 12.0 | 80.6 ± 1.0 | 82.7 ± 0.4 | 270 ± 20 | 1100 | 834 | 715 ± 24 | $A_{10}B_{10}$ | 741 |
| | 13.3 | 66.8 ± 0.1 | 66.9 ± 0.1 | 200 ± 10 | 830 | 504 | 479 ± 20 | $A_8B_8$ | 607 |
| | 14.9 | 59.6 ± 1.5 | 67.2 ± 3.3 | 180 ± 30 | 508 | 266 | 318 ± 27 | $A_4B_4$ | 299 |
| active-short | | | | | | | | | |
| | 13.6 | 50.9 ± 0.6 | 53.2 ± 0.6 | 170 ± 10 | 346 | 208 | 243 ± 13 | $A_4B_4$ | 299 |
| | 14.8 | 39.4 ± 0.1 | 40.3 ± 0.1 | 140 ± 20 | 228 | 153 | 147 ± 9 | $A_2B_2$ | 149 |
| | 16.3 | 34.0 ± 0.4 | 33.9 ± 0.3 | 110 ± 10 | 183 | 118 | 119 ± 6 | $A_2B_2$ | 149 |
| active | | | | | | | | | |
| | 15.4 | 33.9 ± 0.1 | 34.1 ± 0.1 | 100 ± 10 | 201 | 134 | 147 ± 7 | $A_2B_2$ | 149 |

[a]Estimated from the distance value at which the P(r) function calculated from indirect Fourier transform approaches zero; [b]From the Porod volume $V_P$ according to the empirical relation MW~0.625$V_P$ (7); [c]From the Bayesian inference approach based on concentration-independent methods (10).

**Table S3**. Cross correlation values for A-subunit or B-subunit positioned in the contact region between adjacent tetramers in the various GAPDH oligomers. The values have been calculated using the "fit" command as implemented in UCSF Chimera (20).

| | **GAPDH oligomer** | | | | | | | |
|---|---|---|---|---|---|---|---|---|
| | $A_4B_4$ | | $A_8B_8$ Main conf. | | $A_8B_8$ Alt. Conf. | | $A_{10}B_{10}$ | |
| | A-sub. | B-sub. | A-sub. | B-sub. | A-sub. | B-sub. | A-sub. | B-sub |
| | 0.9416 | 0.9464 | 0.9270 | 0.9334 | 0.9434 | 0.9493 | 0.9744 | 0.9759 |
| | 0.9408 | 0.9462 | 0.9270 | 0.9334 | 0.9431 | 0.9493 | 0.9744 | 0.9759 |
| | 0.9405 | 0.9449 | 0.9268 | 0.9334 | 0.9411 | 0.9487 | 0.9743 | 0.9758 |
| | 0.9397 | 0.9447 | 0.9268 | 0.9326 | 0.9411 | 0.9487 | 0.9743 | 0.9758 |
| | | | 0.9261 | 0.9325 | 0.9406 | 0.9464 | 0.9743 | 0.9758 |
| | | | 0.9260 | | 0.9406 | 0.9461 | 0.9743 | 0.9758 |
| | | | | | | 0.9461 | 0.9743 | 0.9758 |
| | | | | | | | | 0.9758 |
| Average | 0.9410 | 0.9460 | 0.9270 | 0.9330 | 0.9420 | 0.9480 | 0.9740 | 0.9760 |
| SD | 0.0008 | 0.0009 | 0.0004 | 0.0005 | 0.0013 | 0.0015 | 0.00005 | 0.00005 |
| t-test | 0.000170926 | | $5.5205 \cdot 10^{-9}$ | | $6.80329 \cdot 10^{-6}$ | | $7.6780 \cdot 10^{-17}$ | |

**Table S4**. GAPDH oligomers average interface areas and ΔG calculated by PDBePISA (22).

| **GAPDH oligomer** | **N° Interfaces** | **Total Interface Area** | **Single Interface Area** | **$\Delta G_{int}$** | **$\Delta G_{diss}$** |
|---|---|---|---|---|---|
| | (#) | (Å$^2$) | | (kcal mol$^{-1}$) | |
| $A_4B_4$ | 1 | 656 | 656 | -234.6 | 35.9 |
| $A_4B_4$ (no CTE) | 1 | 403 | 403 | -212.4 | -12.4 |
| $A_8B_8$ Main Conf. | 4 | 2641 | 660 | -537.9 | 41.0 |
| $A_8B_8$ Main Conf. (no CTE) | 4 | 1795 | 449 | -484.0 | -17.0 |
| $A_8B_8$ Alt. Conf. | 4 | 2501 | 625 | -732.7 | 35.5 |
| $A_8B_8$ Alt. Conf. (no CTE) | 4 | 1684 | 421 | -668.6 | -17.0 |
| $A_{10}B_{10}$ | 5 | 1139 | 228 | -618.6 | -34.7 |

**Table S5**. Results of the optimization of the selected averaged SAXS profiles in the SEC-SAXS experiments as a linear combination of AB-GAPDH oligomers. The $\chi^2$ value obtained by fitting the data with a single structural model are reported in the last column for comparison.

| | | Optimized volume fractions | | | | | Calculated | | | |
|---|---|---|---|---|---|---|---|---|---|---|
| Sample | <$V_{SEC}$> (ml) | AB (OR) | $A_2B_2$[a] | $A_4B_4$ | $A_8B_8$[b] | $A_{10}B_{10}$ | MW (kDa) | $R_g$ (Å) | $\chi^2$ | $\chi^{2c}$ |
| inactive | | | | | | | | | | |
| | 12.0 | - | 0 | 0.154 ± 0.005 | 0.057 ± 0.008 | 0.789 ± 0.006 | 687 | 77.1 | 2.4 | 4.9 |
| | 13.3 | - | 0 | 0.131 ± 0.001 | 0.753 ± 0.042 | 0.116 ± 0.002 | 591 | 66.8 | 4.9 | 19.2 |
| | 14.9 | - | 0.079 ± 0.176 | 0.719 ± 0.242 | 0.162 ± 0.113 | 0.040 ± 0.035 | 357 | 57.8 | 1.0 | 1.0 |
| active-short | | | | | | | | | | |
| | 13.6 | 0 | 0.114 ± 0.053 | 0.849 ± 0.069 | 0.037 ± 0.023 | 0 | 294 | 51.5 | 1.0 | 1.0 |
| | 14.8 | 0.146 ± 0.002 | 0.570 ± 0.002 | 0.284 ± 0.001 | 0 | 0 | 181 | 42.2 | 10.1 | 143 |
| | 16.3 | 0.332 ± 0.015 | 0.650 ± 0.013 | 0.018 ± 0.006 | 0 | 0 | 128 | 33.0 | 0.9 | 1.4 |
| active | | | | | | | | | | |
| | 15.4 | 0.101 ± 0.007 | 0.854 ± 0.006 | 0.045 ± 0.003 | 0 | 0 | 149 | 34.6 | 1.1 | 2.1 |

[a]From the $A_2B_2$ crystal structure (PDB ID 2PKQ) (11); [b]Both the cryoEM derived models of $A_8B_8$ were included (main population and alternative conformation) and here the sum of their volume fractions is reported. Their theoretical scattering profile is almost indistinguishable, as seen in Fig. S9; [c]By fitting the selected data with the theoretical scattering profile of a single structural model with CRYSOL 3.0 in fitting mode, as explained in the caption of Fig. S10.

**Table S6**. Summary of the SC-SAXS data analysis.

| Sample | inactive | | | | | | | |
|---|---|---|---|---|---|---|---|---|
| Concentration (mg·mL$^{-1}$) | 1.89 | 1.52 | 1.18 | 0.67 | 0.52 | 0.39 | 0.2 | 0.08 |
| **Guinier fit** | | | | | | | | |
| $R_g$ (Å) | 82.2 | 84.6 | 82.0 | 81.0 | 80.8 | 81.1 | 80.8 | 88.9 |
| σ($R_g$) (Å) | 3.4 | 4.5 | 8.7 | 17.5 | 20.6 | 60.0 | 16.9 | 11.6 |
| I(0) [kDa] | 530 | 546 | 513 | 561 | 509 | 490 | 470 | 509 |
| σ(I(0)) | 2 | 1 | 1 | 3 | 3 | 4 | 4 | 9 |
| First q point (Å$^{-1}$) | 0.0075 | 0.0075 | 0.0089 | 0.0099 | 0.0100 | 0.0108 | 0.0080 | 0.0075 |
| Last q point (Å$^{-1}$) | 0.015 | 0.012 | 0.015 | 0.016 | 0.016 | 0.016 | 0.016 | 0.015 |
| Auto$R_g$ quality | 0.85 | 0.78 | 0.78 | 0.74 | 0.71 | 0.41 | 0.56 | 0.42 |
| **Indirect Fourier transform** | | | | | | | | |
| $R_g$ (Å) | 86.0 | 85.6 | 86.9 | 88.2 | 86.9 | 88.6 | 85.6 | 93.7 |
| σ($R_g$) (Å) | 0.2 | 0.3 | 0.4 | 0.5 | 0.8 | 1.3 | 1.7 | 2.3 |
| I(0) [kDa] | 536.0 | 543.5 | 521.6 | 581.7 | 525.1 | 510.9 | 477.1 | 511.6 |
| σ(I(0)) | 0.9 | 1.0 | 1.6 | 2.3 | 2.9 | 4.6 | 5.7 | 10.7 |
| VP (·$10^{-3}$ Å$^3$) | 1240 | 1220 | 1260 | 1310 | 1270 | 1370 | 1210 | 1460 |
| First q point (Å$^{-1}$) | 0.0075 | 0.0075 | 0.0089 | 0.0099 | 0.0100 | 0.0108 | 0.0800 | 0.0750 |
| Last q point (Å$^{-1}$) | 0.35 | 0.35 | 0.35 | 0.35 | 0.35 | 0.35 | 0.35 | 0.35 |
| Dmax imposed for P(r) (Å) | 340 | 340 | 340 | 340 | 340 | 340 | 340 | 340 |
| Dmax variability estimate (Å) | 50 | 50 | 50 | 50 | 50 | 50 | 50 | 50 |
| GNOM quality estimate | 0.53 | 0.53 | 0.53 | 0.54 | 0.50 | 0.52 | 0.55 | 0.46 |
| MW($V_P$)[a] (kDa) | 775 | 763 | 788 | 819 | 794 | 856 | 756 | 913 |
| MW ($V_c$)[b] (kDa) | 626 | 627 | 625 | 619 | 608 | 658 | 607 | 573 |
| MW (MoW)[c] (kDa) | 661 | 663 | 658 | 625 | 589 | 687 | 580 | 466 |

| Sample | active | | | | |
|---|---|---|---|---|---|
| Concentration (mg·mL$^{-1}$) | 2 | 1 | 0.5 | 0.25 | 0.1 |
| Guinier fit | | | | | |
| $R_g$ (Å) | 66.5 | 63.0 | 60.5 | 54.5 | 50.2 |
| σ($R_g$) (Å) | 1.5 | 1.0 | 2.9 | 2.8 | 1.5 |
| I(0) [kDa] | 134.1 | 111.6 | 95.0 | 76.5 | 63.7 |
| σ(I(0)) | 0.2 | 0.2 | 0.3 | 0.3 | 0.7 |
| First q point (Å$^{-1}$) | 0.0097 | 0.0055 | 0.0060 | 0.0060 | 0.0070 |
| Last q point (Å$^{-1}$) | 0.0187 | 0.0192 | 0.0187 | 0.0234 | 0.0258 |
| Auto$R_g$ quality | 0.75 | 0.64 | 0.82 | 0.45 | 0.26 |
| **Indirect Fourier transform** | | | | | |
| $R_g$ (Å) | 70.5 | 66.6 | 63.7 | 59.7 | 53.9 |
| σ($R_g$) (Å) | 0.3 | 0.4 | 0.6 | 1.1 | 2.1 |
| I(0) [kDa] | 137.0 | 113.0 | 95.8 | 78.2 | 64.4 |

| | | | | | |
|---|---|---|---|---|---|
| σ(I(0)) | 0.3 | 0.2 | 0.4 | 0.6 | 1.1 |
| $V_P$ ($\cdot 10^{-3}$ Å$^3$) | 502 | 431 | 363 | 329 | 258 |
| First q point (Å$^{-1}$) | 0.0097 | 0.0055 | 0.0060 | 0.0060 | 0.0070 |
| Last q point (Å$^{-1}$) | 0.35 | 0.35 | 0.35 | 0.35 | 0.35 |
| $D_{max}$ imposed for P(r) (Å) | 300 | 280 | 250 | 240 | 230 |
| $D_{max}$ variability estimate (Å) | 100 | 20 | 50 | 30 | 30 |
| GNOM quality estimate | 0.48 | 0.52 | 0.46 | 0.47 | 0.42 |
| MW($V_P$)[a] (kDa) | 314 | 270 | 227 | 206 | 161 |
| MW ($V_c$)[b] (kDa) | 312 | 253 | 185 | 168 | 139 |
| MW (MoW)[c] (kDa) | 377 | 329 | 219 | 217 | 179 |

[a]From the Porod volume $V_P$ according to the empirical relation MW~0.625$V_P$ (7); [b]From the volume of correlation $V_c$ ($q_{max}$ for integration 0.25 Å$^{-1}$) (8); [c]From the Porod invariant ($q_{max}$ for integration 0.25 Å$^{-1}$) (9).

**Table S7**. Results of the optimization of the SC-SAXS data for concentration series of active and inactive AB-GAPDH samples as a linear combination of AB-GAPDH oligomers.

| Sample | c (mg·mL$^{-1}$) | Optimized volume fractions | | | | $\chi^2$ | Calculated | |
|---|---|---|---|---|---|---|---|---|
| | | $A_2B_2$ | $A_4B_4$ | $A_8B_8$[a] | $A_{10}B_{10}$ | | MW (kDa) | $R_g$ (Å) |
| inactive | 1.89 | 0 | 0.163 ± 0.001 | 0.377 ± 0.002 | 0.460 ± 0.002 | 19.1 | 634 | 72.6 |
| | 1.52 | 0 | 0.160 ± 0.002 | 0.377 ± 0.003 | 0.463 ± 0.002 | 13.1 | 635 | 72.6 |
| | 1.18 | 0 | 0.162 ± 0.002 | 0.362 ± 0.003 | 0.476 ± 0.003 | 8.8 | 636 | 72.9 |
| | 0.67 | 0 | 0.161 ± 0.003 | 0.334 ± 0.005 | 0.504 ± 0.004 | 4.8 | 641 | 73.3 |
| | 0.52 | 0 | 0.157 ± 0.004 | 0.329 ± 0.006 | 0.513 ± 0.005 | 3.3 | 644 | 73.4 |
| | 0.39 | 0 | 0.147 ± 0.005 | 0.307 ± 0.008 | 0.545 ± 0.007 | 2.0 | 652 | 73.9 |
| | 0.20 | 0 | 0.146 ± 0.009 | 0.372 ± 0.016 | 0.482 ± 0.012 | 1.0 | 643 | 73.0 |
| | 0.08 | 0 | 0.128 ± 0.023 | 0.355 ± 0.037 | 0.517 ± 0.028 | 0.8 | 653 | 73.5 |
| active | | | | | | | | |
| | 2.00 | 0.181 ± 0.011 | 0.507 ± 0.017 | 0.042 ± 0.009 | 0.270 ± 0.003 | 11.4 | 412 | 66.8 |
| | 1.00 | 0.335 ± 0.021 | 0.445 ± 0.031 | 0.022 ± 0.016 | 0.197 ± 0.005 | 2.2 | 350 | 63.5 |
| | 0.50 | 0.453 ± 0.008 | 0.395 ± 0.009 | 0 | 0.152 ± 0.003 | 1.7 | 305 | 60.5 |
| | 0.25 | 0.551 ± 0.016 | 0.336 ± 0.019 | 0 | 0.113 ± 0.005 | 0.8 | 272 | 57.3 |
| | 0.10 | 0.643 ± 0.041 | 0.283 ± 0.046 | 0 | 0.074 ± 0.013 | 0.8 | 240 | 53.1 |

[a]Both the cryoEM derived models of $A_8B_8$ were included (main population and alternative conformation) and here the sum of their volume fractions is reported. Their theoretical scattering profile is almost indistinguishable, as seen in Fig. S9.

**Table S8.** Summary of SAXS data acquisition information, sample details, and data analysis software used.

| (A) Sample details for the SEC-SAXS experiments | | | |
|---|---|---|---|
| | inactive | active-short | active |
| Loading concentration (mg·mL$^{-1}$) | 13 | 11 | < 5 |
| Injection volume (μL) | 100 | 200 | 200 |
| Storage buffer composition | 25 mM K-phosphate, pH 7.5, 0.1 mM NAD$^+$ | 25 mM K-phosphate, pH 7.5, 5 mM reduced DTT, 20 mM NADP$^+$ 1,3-bisphosphoglirate* | 25 mM K-phosphate, pH 7.5, 5 mM reduced DTT, 20 mM NADP$^+$ 1,3-bisphosphoglirate* |
| Elution buffer composition | 25 mM K-phosphate, pH 7.5, 0.1 mM NAD$^+$ | 25 mM K-phosphate, pH 7.5, 0.1 mM NADP$^+$ | 25 mM K-phosphate, pH 7.5, 0.1 mM NADP$^+$ |

*obtained by incubation of phosphoglycerate kinase, 20 U ml$^{-1}$, with 15 mM 3-phosphoglyceric acid, 10 mM ATP and 5 mM $MgCl_2$

| (B) SAXS data collection parameters for the SEC-SAXS experiments | |
|---|---|
| Source, instrument | ESRF, BM29 (23) |
| Wavelength (Å) | 0.9919 |
| Sample-to-detector distance (m) | 2.872 |
| $q=4\pi\sin(\theta)/\lambda$ ($2\theta$ scattering angle) range (Å$^{-1}$) | 0.005-0.45 |
| Absolute scaling method | water scattering I(0)= 0.01632 cm$^{-1}$, protein partial specific volume 0.735 cm$^3$ g$^{-1}$ |
| Exposure time (s) | 1 |
| Capillary path length (mm) | 1.8 |
| SEC column | Superdex 200 10/300 GL (GE Healthcare) |
| Flow rate (mL·min$^{-1}$) | 0.5 |
| SEC column temperature (ºC) | 22 |

| (C) Sample details for the SC-SAXS experiments | | |
|---|---|---|
| | inactive | active |
| Concentration range (mg·mL$^{-1}$) | 0.08-1.89 | 0.1-2.0 |
| Storage and dilution buffer composition | 25 mM K-phosphate, pH 7.5, 1 mM NAD$^+$ | 25 mM K-phosphate, pH 7.9, 1 mM NADP$^+$ |

| (D) SAXS data collection parameters for the SC-SAXS experiments | | |
|---|---|---|
| | inactive | active |
| Source, instrument | ESRF, BM29 (23) | |
| Wavelength (Å) | 0.9919 | |
| sample-to-detector distance (m) | 2.872 | 2.864 |
| q-measurement range (Å$^{-1}$) | 0.005-0.45 | |
| Absolute scaling method | water scattering I(0)= 0.01632 cm$^{-1}$, protein partial specific volume 0.735 cm$^3$ ·g$^{-1}$ | |
| Capillary path length (mm) | 1.8 | |
| Injection volume (μL) | 50 | 60 |
| Exposure time (s) | 1 | 2 |
| Number of exposures | 10 | 10 |
| Extra flow time (s) | 10 | 10 |
| Sample temperature (ºC) | 4 | 5 |

| (E) Software employed for SAS data reduction, analysis, and interpretation | |
|---|---|
| Solvent subtraction, averaging and basic analysis (Guinier fit, P(r), Porod Volume) | Matlab scripts, ATSAS 2.8 (4) |
| Theoretical intensity calculations | CRYSOL 3.0, OLIGOMER |
| Molecular graphics | PyMOL 1.8 |

**Table S9**. CryoEM data collection and data processing parameters.

| | |
|---|---|
| **DATA COLLECTION** | |
| Microscope model | Thermo Fisher Scientific Tecnai Polara F30 |
| Detector type | GATAN K2 Summit |
| Imaging mode | Bright field |
| Accelerating voltage, kV | 300 |
| Nominal/Calibrated magnification | 31000 |
| Pixel size, Å | 1.21 |
| Total exposure time, sec | 4 |
| Total Number of collected stacks | 2228 |
| Number of stacks used in the analysis | 1988 |
| Total dose per stack, $e^{-}/Å^2$ | 42 |
| Number of frames per stack | 40 |
| Defocus range, μm | from -1.5 to -3.5 |
| Defocus step, μm | 0.15 |
| **DATA PROCESSING, GLOBAL RESOLUTION (Å) AND EMDB ID** | |
| 3D reconstruction software package | Relion 3.0 |
| $A_2B_2$ | |
| Extracted particles | 48558 |
| Refined particles | 19636 |
| Symmetry | D2 |
| FSC0.143 (unmasked/masked) | 6.5/6.3 |
| Local resolution range, Å | 3.7-9.7 |
| EMBD ID | 13824 |
| $A_4B_4$ | |
| Extracted particles | 31023 |
| Refined particles | 20777 |
| Symmetry | C1 |
| FSC0.143 (unmasked/masked) | 13.1/8.9 |
| Local resolution range, Å | 4-15 |
| EMBD ID | 13825 |
| $A_8B_8$ main conformer | |
| Extracted particles | 64130 |
| Refined particles | 23611 |
| Symmetry | C2 |
| FSC0.143 (unmasked/masked) | 7.4/5.7 |
| Local resolution range, Å | 3.7-10.2 |
| EMBD ID | 13826 |
| $A_8B_8$ alternative conformer | |
| Extracted particles | 64130 |
| Refined particles | 10768 |
| Symmetry | C2 |
| FSC0.143 (unmasked/masked) | 8.2/7.1 |
| Local resolution range, Å | 4-11.5 |
| EMBD ID | 13827 |
| $A_{10}B_{10}$ | |

| | |
|---|---|
| Total extracted particles | 33067 |
| Refined particles | 7352 |
| Symmetry | C5 |
| FSC0.143 (unmasked/masked) | 15.1/13 |
| Local resolution range, Å | 4.7-14.7 |
| EMBD ID | 13828 |
